# Alternate RNA decoding results in stable and abundant proteins in mammals

**DOI:** 10.1101/2024.08.26.609665

**Authors:** Shira Tsour, Rainer Machne, Andrew Leduc, Simon Widmer, Eunice Koo, Jeremy Guez, Konrad Karczewski, Nikolai Slavov

**Author notes:** Data & code: decode.slavovlab.net.

## Abstract

Amino acid substitutions may substantially alter protein stability and function, but the contribution of substitutions arising from alternate translation (deviations from the genetic code) is unknown. To explore it, we analyzed deep proteomic and transcriptomic data from over 1,000 human samples, including 6 cancer types and 26 healthy human tissues. This global analysis identified 60,803 fragmentation spectra corresponding to 8,801 unique substitution sites in proteins derived from 1,782 genes, including 2,000 confidently localized sites. Some substitutions are shared across samples, while others exhibit strong tissue-type and cancer specificity. Surprisingly, products of alternate translation are more abundant than their canonical counterparts for hundreds of proteins, suggesting sense codon recoding. Recoded proteins include transcription factors, proteases, signaling proteins, and proteins associated with neurodegeneration. Mechanisms contributing to substitution abundance include protein stability, codon frequency, codon-anticodon mismatches, and RNA modifications. We characterize how alternatively translated proteoform ratios vary across protein domains, tissue types and cancers. The substitution ratios are positively associated with intrinsically disordered regions and genetic polymorphisms in gnomAD, though the polymorphisms cannot account for the substitutions. The sequence, relative abundance, and the tissue-specificity of alternatively translated proteins are conserved between human and mouse. These results demonstrate the contribution of alternate translation to diversifying mammalian proteomes, and its association with protein stability, tissue-specific proteomes, and diseases.

## Introduction

Genetic mutations, RNA editing, and “alternate translation” (mRNA translation deviating from the genetic code) may result in amino acid substitutions. Some substitutions arising from mutations profoundly change protein activity. For example, the V600E phosphomimetic substitution in BRAF destabilizes hydrophobic interactions and constitutively increases BRAF activity up to 500-fold, inducing tumorigenesis^1^. Similarly, the H1047R substitution in the catalytic subunit *p*110*α* of PI3K causes extensive cellular remodeling^2^. Substitutions introduced by RNA editing also alter protein functions, such as the A-to-I editing of the glutamate receptor mRNA^3^.

In the absence of genetic mutations and RNA editing detectable in RNA sequences, amino acid substitutions may arise from translation alternate to the genetic code. Such alternate decoding can occur when translating all open reading frames (short or long, annotated or not annotated as protein-coding), but it remains less characterized. For decades, amino acid substitutions were detected based on gel shifts of radioactively labeled proteins^4^. Mass spectrometry (MS) greatly increased the power of detecting substitutions^5,6^, and deep learning predictions of peptide elution times and MS fragmentation spectra are facilitating the validation of non-canonical amino acid sequences^7–9^. Translational substitutions are mostly studied in the context of amino acid starvation and antibiotic treatments as errors^10–12^ but have also been detected in healthy organisms, mostly unicellular microorganisms, where the prevalence and the stability of alternatively translated proteins are presumed low^12–14^. They are often considered as translation errors resulting in low level molecular noise, though studies have suggested that non-cognate amino acid acylation may confer beneficial functions^15,16^. Another possible mechanism may involve mRNA modifications, such as pseudouridylation, which may recode stop codons to promote readthrough^17,18^, though its impact on the sequence of endogenous proteins is uncharacterized.

If the rate of alternate translation is low, the abundance and significance of its protein products may be low as well. However, the abundance of proteoforms harboring amino acid substitutions is not determined solely by the rate of their synthesis. It also depends on the rate of their degradation. While most substitutions are likely to destabilize the substituted proteoform, some might stabilize them; if stabilized by substitution, a proteoform may accumulate to high abundance. Whether such protein stabilization contributes to elevated levels of proteins synthesized by alternate RNA decoding remains unknown. To explore this question, we systematically identified and quantified amino acid substitutions across healthy and cancer human and mouse tissues. We identified, validated and characterized hundreds of abundant amino acid substitutions, including mechanisms contributing to their origin and their impact on protein stability.

## Results

### Systematic identification and validation of amino acid substitutions

To globally identify peptide modifications, we analyzed paired MS and RNA-seq data from 1,094 human samples described in Extended Data Fig. 1a,b. They included 6 cancer types (renal, uterine, pancreatic, breast, prostate, lung squamous cell carcinoma and lung adenocarcinoma) and matched normal adjacent tissue from the Clinical Proteomic Tumor Analysis Consortium (CPTAC)^20–25^ and 26 tissue types from healthy individuals^26^, Fig. 1a. We included TMT-labeled and label-free MS data from different laboratories to explore peptide modifications across tissue types and disease states independent from biases specific to particular datasets. To minimize the influence of technical variation leading to missing data (peptides identified only in some samples), our analysis focuses on abundance ratios between quantified peptides, which are less affected by variations in data acquisition parameters.

**Figure 1.**
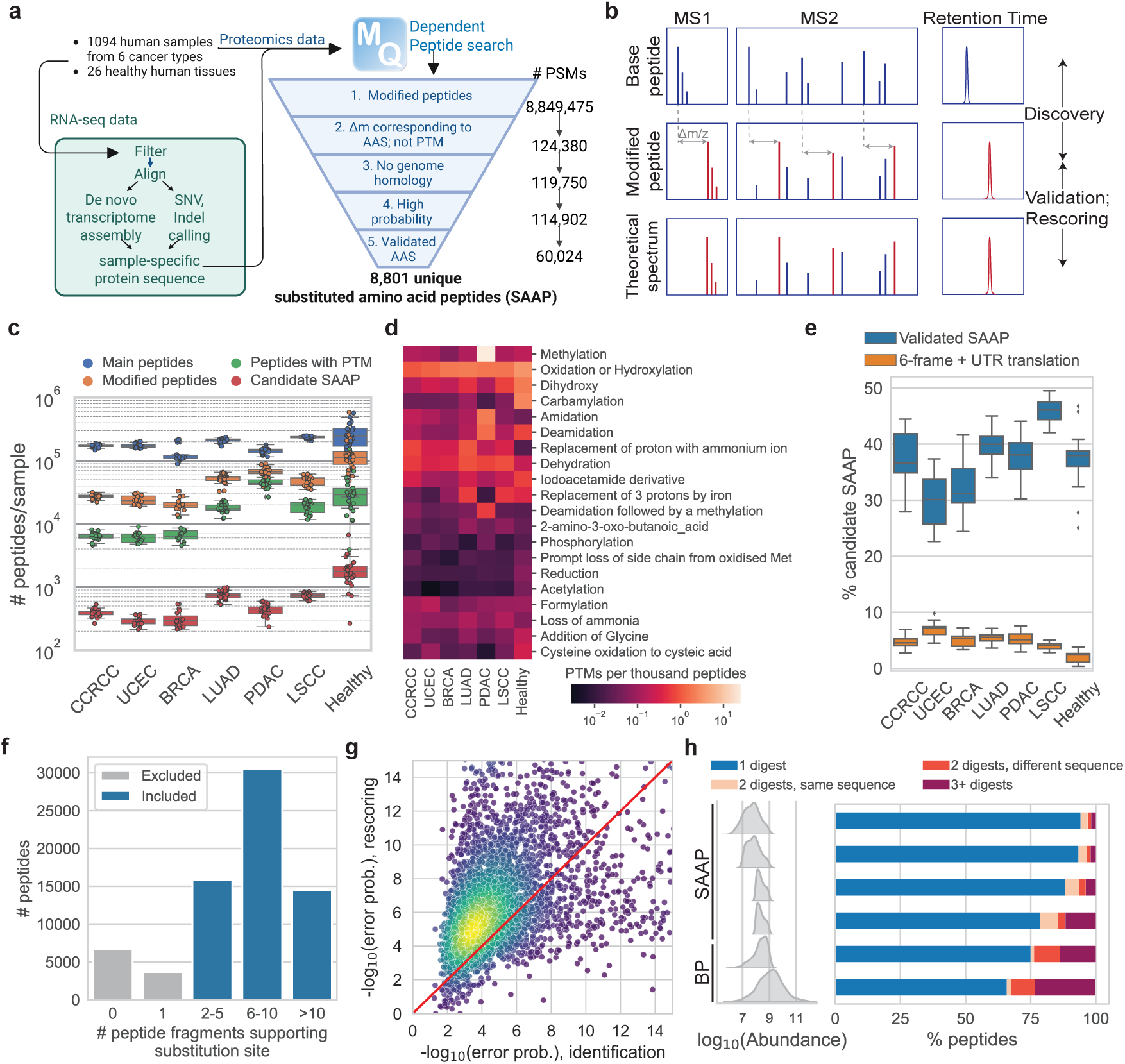
Identification and validation of amino acid substitutions. (**a**) A schematic of the data processing workflow. RNA-seq data is *in-silico* translated to patient-specific protein databases, which are used to search patient-matched MS proteomics data for peptides with modifications. The resulting peptides are filtered and further validated. (**b**) Search strategy for discovering modified peptides in MS spectra with ModifiComb. Spectra with systematic mass shifts of the precursor and fragment ions from identified (base) peptides correspond to modified peptides^5^. (**c**) Number of database peptides (blue), modified peptides (any mass shift, orange), peptides with PTMs (mass shift of a known modification, green) or candidate SAAP (mass shift of a substitution, red). Distributions are across CPTAC datasets or across tissue samples (label-free data). (**d**) The number of PTM peptides identified per 1,000 unmodified peptides for the most common PTMs. (**e**) Percentages of candidate SAAP that are discarded as potential products of 6-frame translation (orange) or validated by database-based search and used in further analysis (blue). (**f**) Distribution of the number of detected fragment ions providing evidence for each substitution. Only substitutions supported by 2 or more fragment ions were analyzed further. (**g**) Re-scoring of validated SAAP. Confidence of peptide sequence identification is higher after re-scoring with Oktoberfest^8^ (y-axis) than initially determined in the validation search with Andromeda^19^ (x-axis). (**h**) Identification of AAS in tonsil proteomes from multiple enzymatic digests. Percentage of SAAP validated in 1 (blue), 2 (orange) or 3 or more (purple) digests, binned by SAAP abundance. The percentage of BP observed in multiple digests is also shown as a control.

To identify modified peptides, we first predicted a protein sequence database for each patient sample by using the genetic code to *in-silico* translate the corresponding transcriptome. Transcriptomes were assembled from paired-end Illumina reads (minimum of 120 million per cancer sample, 18 million per healthy tissue) using a workflow adapted from Galaxy^27^, Fig. 1a. The mean sequence coverage for all transcripts exceeds 98% (median 100%), corresponding to about 71,000 transcripts per sample with 100% sequence coverage, Extended Data Fig. 1c,d. Using the patient-specific databases, we search the MS data with the dependent peptide algorithm of MaxQuant^6^, which implements ModifiComb developed by Savitski *et al.*^5^. This algorithm tests whether MS2 spectra not matched to a database sequence correspond to modifications of peptides identified in the sample. Such correspondence is reflected in systematic mass shifts in the precursor peptide ions and the associated peptide fragments^5^, Fig. 1b, and has been used previously to detect substitutions in bacteria and yeast^13^.

Our dependent peptide search identified almost 9 million spectra matched to modified peptides, 40% of which have mass shifts corresponding to 412 distinct known post-translational modifications (PTMs), Fig. 1c,d, Extended Data Fig. 1e; 124,000 of these peptides exhibited mass shifts consistent with 255 unique amino acid substitutions and no other known modifications, Fig. 1c, Extended Data Fig. 1f. The occurrence of known PTMs with biological and technical origins, such as methionine oxidation, deamidation, and carbamylation, is similar across cancer types, tissue types and datasets, Fig. 1d. These PTMs are reported in Supplemental Data Table 1 and can support much further analysis, which is beyond the scope of this article. Deeper datasets, such as the healthy tissues, have more identified peptides of all types, Fig. 1c.

### Sequence confirmation of amino acid substitutions

We focused on substituted amino acid peptides (which we call SAAP) supported by good RNA sequence coverage and peptide fragmentation spectra, Supplemental Fig. 1, 2 and 3; Supplemental Data Table 2. Mutations in the genomes of LSCC patients did not correspond to any of the identified AAS, further bolstering their non-genetic origin and the reliability of our transcriptome reconstructions. We systematically applied rigorous filters to minimize false positives, including: (1) removed all peptides with a mass shift consistent with another known PTM; (2) validated SAAP discovered by dependent peptide search with a standard database-search, Fig. 1b,e; (3) required that at least 2 peptide fragments reflect a mass shift corresponding to a localized substitution, Fig. 1f, Extended Data Fig. 1h, Supplemental Fig. 4; (4) validated SAAP by comparing their retention time and fragmentation patterns to deep learning predictions, Fig. 1g, from the re-scoring pipeline Oktoberfest^8^, (5) excluded sequences that could be generated by a 6 reading frame translation of any transcript, Fig. 1e, (6) filtered SAAP at 1% FDR computed only across SAAP, Extended Data Fig. 1i, and (7) removed SAAP corresponding to substitutions at trypsin cleavage sites (K, R); across all datasets, only 24 SAAP had K and R substitutions, and about 90% of them represent peptides with missed cleavages or with substitutions from *K* → *R* or vice versa (Extended Data Fig. 1j), supporting the validity of the results. Our confidence in identified SAAP is bolstered by the validation with standard database-search, Fig. 1e, low mass errors for SAAP, Extended Data Fig. 1g, increased confidence after re-scoring, Fig. 1f, Extended Data Fig. 1i, identification of the same substitution site in a human tissue digested by multiple proteases, Fig. 1h, and accurate retention time prediction, as determined by DeepRTplus^28^, Extended Data Fig. 1k. The database search validated about 40% of the SAAP discovered by DP search, Fig. 1e; this result is expected since the database search does not use the spectra of the main peptides and thus has lower sensitivity. These SAAP were further filtered to remove peptides mapping to immunoglobulins and trypsin, since we could not be confident that these are true instances of alternate translation. For about half of all spectra matched to SAAP, the measured mass shift indicating a substitution is localized with high confidence to a single position (Supplemental Fig. 1, 2 and 4a), and thus cannot arise from multiple combinatorial modifications on different residues adding up to the mass shift; for the remaining SAAP, unknown combinatorial modifications cannot be rigorously disproved (see Methods for details). However, specific hypotheses for explaining observed mass shifts (such as *Q* → *G* versus cleavage of terminal A) may be discriminated based on their fragment ion intensities. The empirical intensities are closer to those predicted for *Q* → *G*, thus suggesting it is the more likely hypothesis, Supplemental Fig. 4c-f. Our extensive filtering likely removes many true SAAP, but it helps establish a robust set of nearly 9,000 unique SAAP - BP pairs, Supplemental Data Table 2. For SAAP with imprecise mass shift localization, we cannot exclude all alternative hypotheses (involving multiple modifications), and thus we establish a set of 2,000 AAS (unique pairs of BP - SAAP) with high localization probability (*>* 0.9) of the substitution, reported in Supplemental Data Table 8. These SAAP are strongly supported by the data and support exploration of the biological implications of all putative alternate translation sites. Blasting all substituted peptides against the 131,000 Ensembl proteins or Uniprot showed that only 110 of 8,948 SAAP sequences have been previously predicted from genetic code translations.

**Figure 2.**
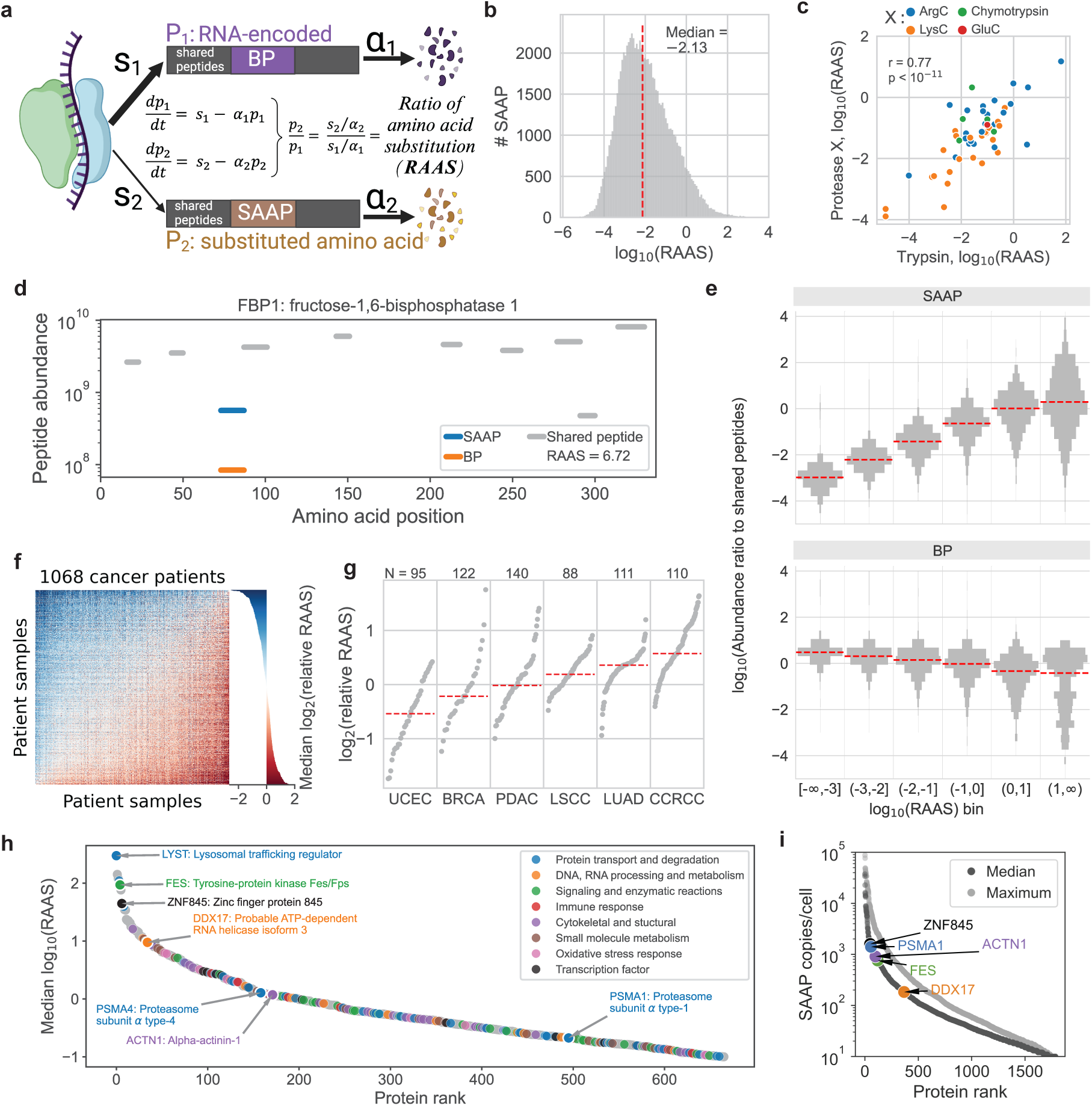
Abundance of proteins with amino acid substitutions. (**a**) Model for the abundance of the canonical (P1) and the alternatively translated (P2) proteoforms. P1 and P2 have base (BP) and substituted (SAAP) peptides, respectively, and shared peptides are common to both. P1/P2 is estimated with the SAAP/BP ratio.(**b**) Distribution of substitution ratios computed for each patient sample in 7 datasets. (**c**) Reproducibility of RAAS for substitutions quantified in multiple digests. Data points are individual SAAP. (**d**) An example protein with a highly abundant substitution showing that the shared peptide abundances are in closer agreement with the SAAP abundance than the base peptide abundance. (**e**) The abundance of shared peptides follows the abundance trends for BP and SAAP predicted from our model (panel a) and the corresponding ratios denoted on the x-axis. (**f**) RAAS fold changes (relative RAAS) were computed for each SAAP identified in a pair of patients, and their medians displayed. The matrix is sorted by column and row means, and the barplot shows median relative RAAS for each row, corresponding to a patient. (**g**) Mean of relative RAAS computed between a patient relative to all other patients, as in (**c**), weighted by number of shared SAAP. N indicates number of patients in each dataset. (**h**) Rank sorted proteins with RAAS *>* 0.1. Proteinsfrom functional groups that are significantly enriched for high RAAS are highlighted. (**i**) Copy number estimates of SAAP by the histone ruler method^32^.

**Figure 3.**
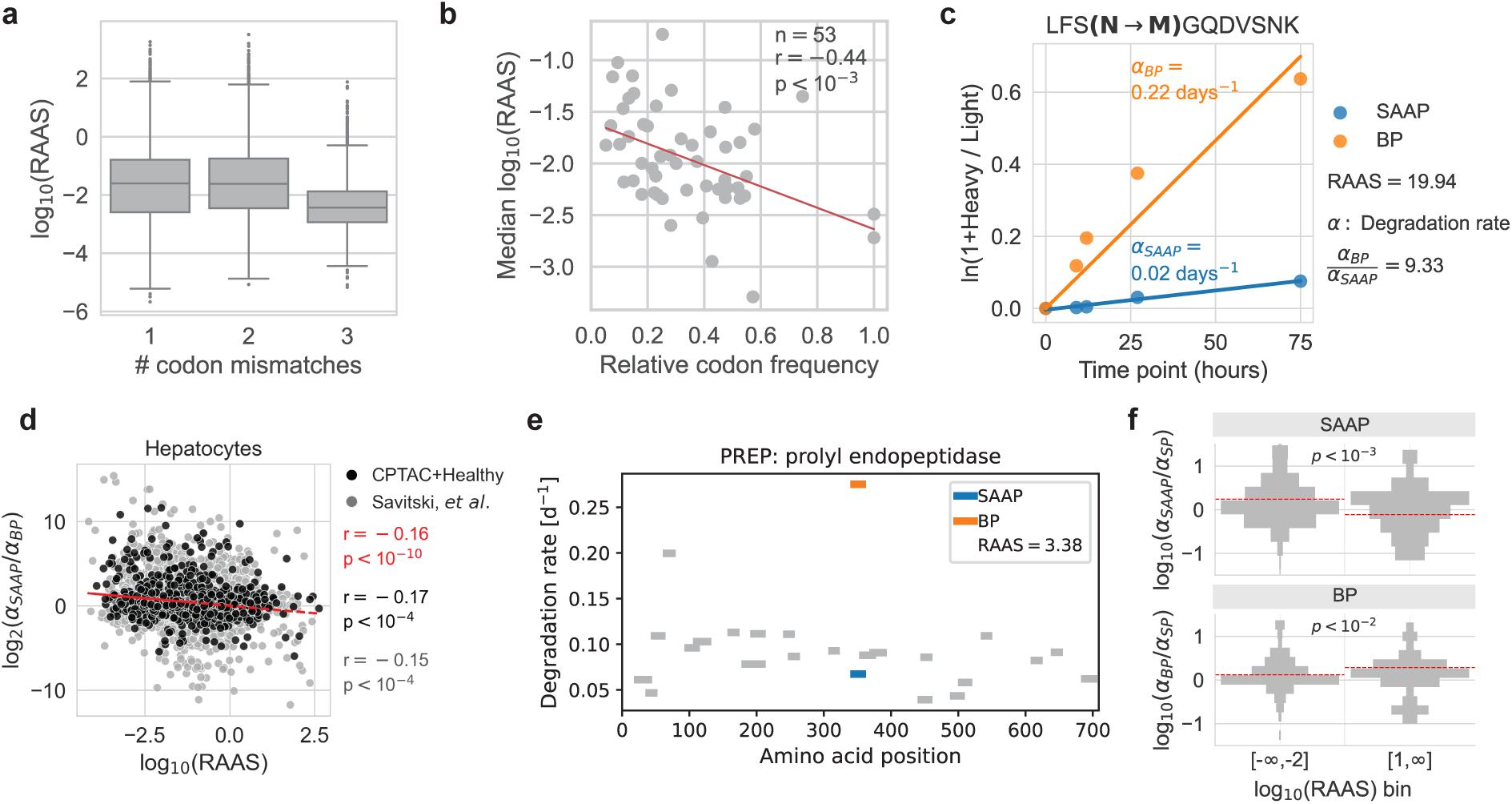
Substitution ratios depend on RNA codons, modifications and proteoform degradation rates. (**a**) RAAS as a function of the minimum number of codon-anticodon mismatches needed for incorporating the detected amino acid. (**b**) The median RAAS of all substitutions mapping to a codon is negatively correlated to the relative frequency of the codon. Least squares fit shown in red, Pearson correlation and associated p-value annotated in plot. See Extended Data Fig. 4 for details. (**c**) Protein degradation rates (*α*) are computed as the slope of *ln*(1 + *h/l*) = *αt*, where *h/l* is the heavy-to-light peptide ratio. For some peptides with abundant substitutions, SAAP degradation rates are up to ten times slower than the degradation rate of the corresponding BP. (**d**) The ratio of degradation rates for SAAP relative to BP is inversely proportional to their RAAS in primary hepatocytes. Substitutions identified only in the hepatocytes are in gray, those also identified in the CPTAC and healthy human tissues in black, and the Pearson correlation for the union is in red. (**e**) Demonstration of a protein with a highly abundant substitution, where the shared peptide degradation rate in closer agreement with the substituted peptide degradation rate than the base peptide degradation rate. (**f**) Degradation rates of substituted peptides are generally faster than degradation rates of shared peptides for proteins with lowly abundant substitution proteoforms. Conversely, degradation rates of base peptides are faster than degradation rates of shared peptides for proteins with highly abundant substitution proteoforms.

**Figure 4.**
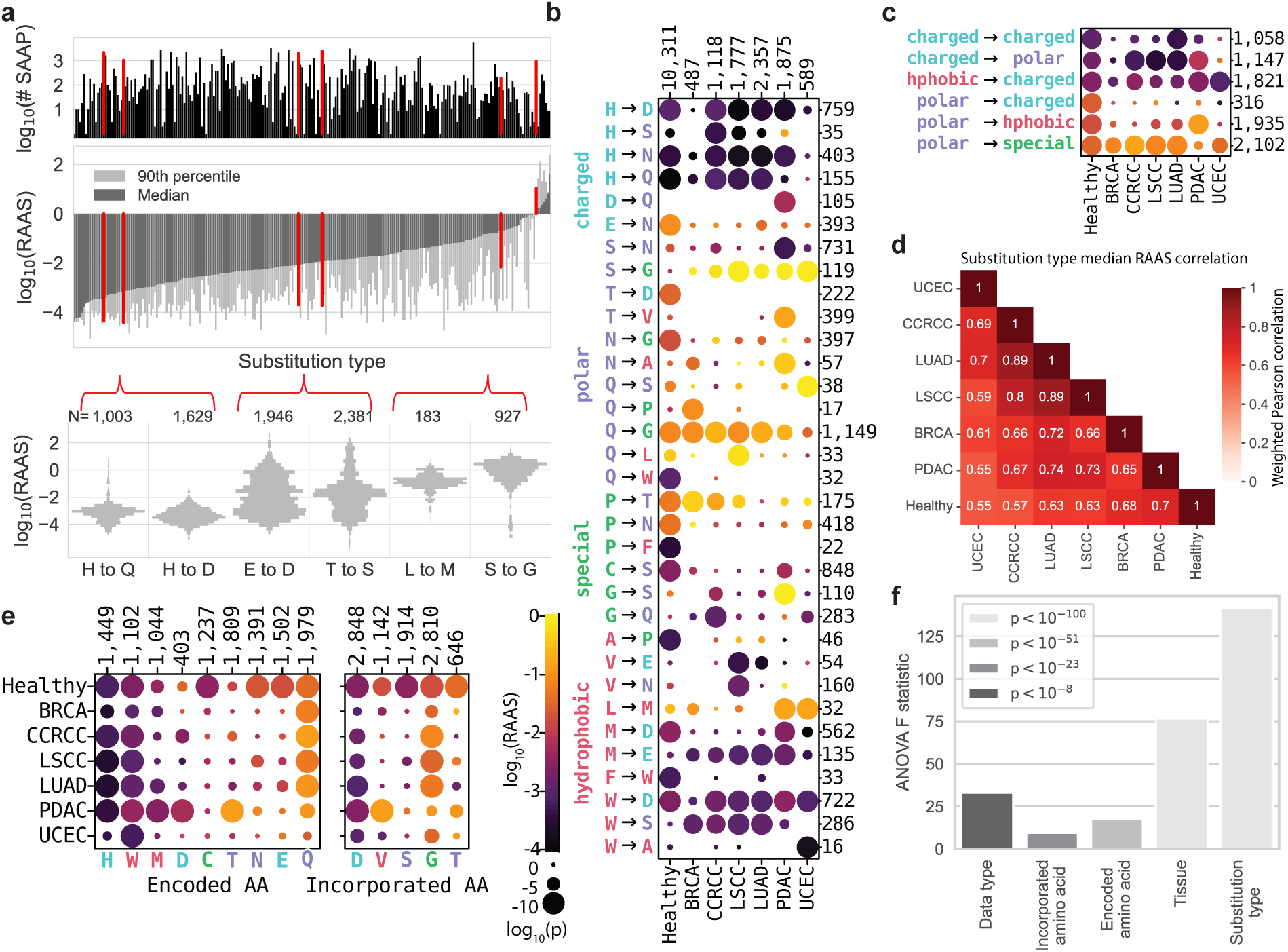
Substitution ratios depend on amino acid and tissue types. (**a**) Number of SAAP (upper panel) and median and 90th percentile RAAS (middle panel) displayed for each substitution type across all datasets. RAAS distributions across all samples are displayed for 6 substitution types (lower panel) spanning the range of RAAS medians, highlighting the variability in RAAS across individual samples, even for the same substitution type. Substitution types displayed were chosen due to large number of data points. (**b**) RAAS dotplot for all amino acid substitution types that had a significantly low or high RAAS distribution (*p* ≤ 10*^−^*^10^) in at least one dataset (columns). The y-axis is grouped by chemical properties of the encoded amino acid. (**c**) RAAS dotplot as in (**b**) but by chemical properties of the encoded and incorporated amino acid. (**d**) Median RAAS profiles across all substitution types are strongly correlated across all dataset pairs (Pearson correlation, weighted by number of SAAP). (**e**) RAAS dotplot as in (**b**) for encoded and incorporated amino acids. (**f**) Results from ANOVA confirming variance in RAAS profiles is driven primarily by substitution type and secondarily by tissue type.

To acquire further evidence for the substituted peptides, we analyzed IP-MS experiments for bait proteins that produced highly confident SAAP, as the reduced proteomic complexity should reduce false positive identifications. We selected 25 bait proteins, representing 56 potential SAAP, pulled from HEK235T cell lines^29^. We then searched each sample for all SAAP sequences from Supplemental Data Table 2. In total we identified 112 unique SAAP across the 25 runs, Supplemental Fig. 5a. Of these SAAP, 34 mapped to the bait protein that was pulled down in the respective experiment, resulting in discovery of SAAP from 18 of the 25 pull-downs, Supplemental Data Table 11. Several potential reasons exist for not identifying SAAP from each bait protein. For example, AAS might affect protein confirmations and thus the interaction between the bait protein and antibody used for the experiment. Additionally, HEK235T may not produce the same SAAP as identified from primary issues.

**Figure 5.**
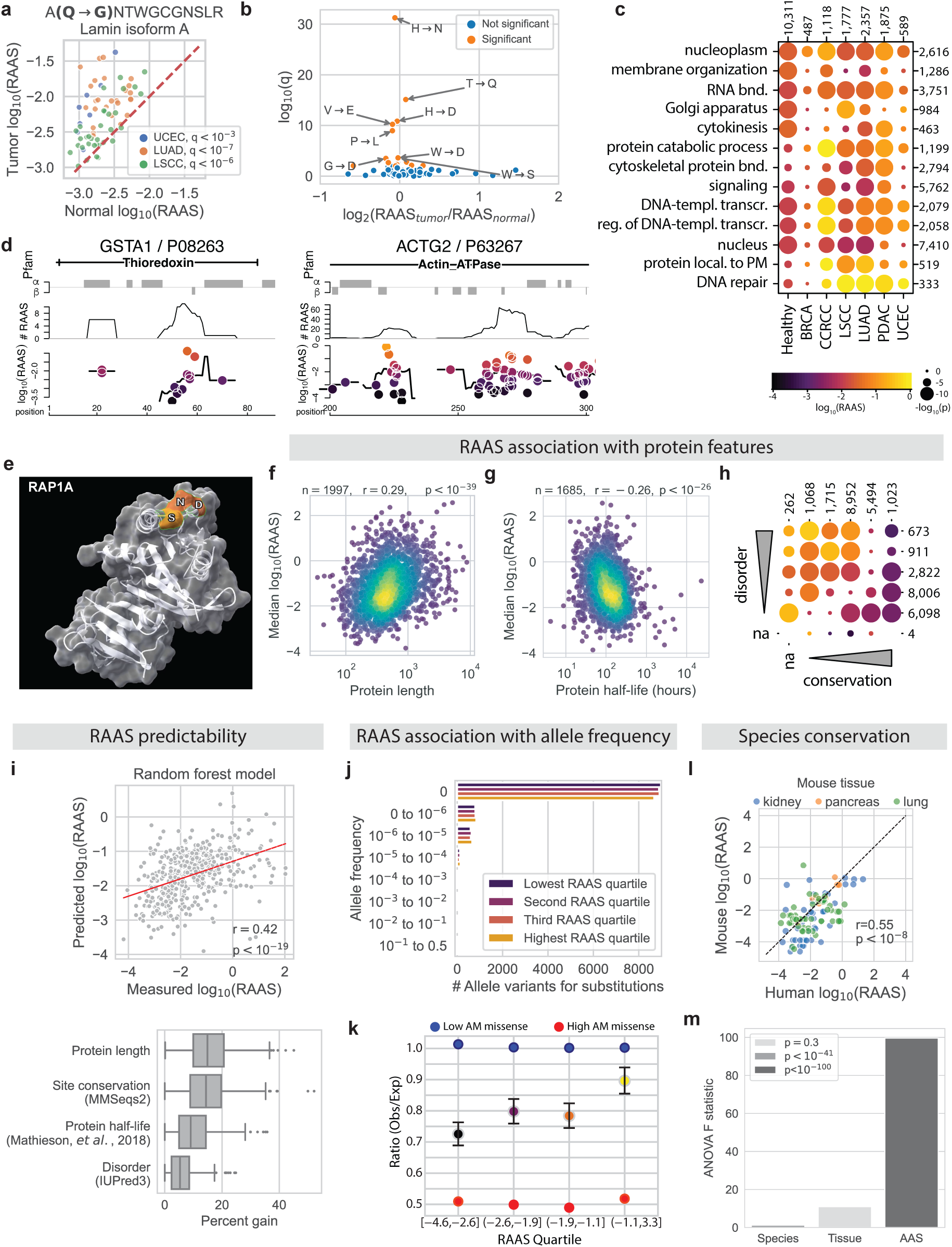
Sequential, structural and functional context of amino acid substitutions. (Caption continued on the next page) (Continued) (**a**) An example SAAP that is significantly more abundant in tumor samples than in normal adjacent tissue from the same patient in 3 cancer types. (**b**) Volcano plot with substitution types that have significantly different RAAS in tumor than in matched normal samples. (**c**) RAAS dotplots for all SAAP in proteins annotated by GO terms with at least one significantly high RAAS distribution (*p* ≤ 10*^−^*^15^). Individual proteins with high RAAS from each GO group are shown in Extended Data Fig. 7. (**d**) Clusters of substitutions in the 1D sequence of a thioredoxin domain (left) and an actin ATPase domain (right). Pfam annotations are shown as black lines, and predicted secondary structures (S4pred), *α*-helices and *β*-sheets, are shown with gray bars. (**e**) A set of three substitutions distant in the 1D sequences but clustered together in the 3D structure of the Ras-related protein, RAP1A. The substitutions have high RAAS, reflected in their color-coding (as in the legend in (**c**)). (**f**) Protein RAAS is significantly positively correlated to protein length. Data points are proteins with substitutions. Protein RAAS is computed as the median RAAS across all SAAP identified in the protein. (**g**) Protein RAAS is significantly negatively correlated to protein half-lives from ref.^39^. Data points are proteins with substitutions. (**h**) RAAS dotplots (as in Fig. 4c) for substitution sites grouped by bins of a predicted disorder score (rows, based in an IUpred3 prediction) and sequence conservation (columns, based on MMseqs2 alignments). (**i**) An XGboost model using protein features described in **(g-h**) is able to accurately predict median RAAS per protein (top panel). Each feature used increases the model’s predictive power (bottom panel). An average split of the data into test and train had a predictive Pearson correlation of 0.44. Additional details can be found in Methods. (**j**) Allele frequency in the gnomAD database for all possible missense variants in alternatively translated codons. (**k**) Observed / expected ratios for all missense variants in alternatively translated codons, determined from analysis of gnomAD database (see Methods). Missense variants are less constrained with increasing RAAS (*p <* 10*^−^*^4^). Data point colors correspond to RAAS quartiles as in (**j**). (**l**) 55 SAAP sequences from human tissues were found to be conserved in mouse lung, kidney and pancreas tissues with significantly correlated RAAS values. (**m**) Substitution type is the most significant driver of variability in RAAS values across the SAAP plotted in (**l**), followed by species and tissue type (ANOVA).

To validate that additional SAAP came from prey proteins in the pull down experiments and not spurious identifications, we plotted the number of non-SAAP peptides shared by the proteins from which each SAAP was identified, Supplemental Fig. 5b. This number was larger on average for SAAP identified from the bait protein, and all SAAP had at least 2 non-SAAP peptides (median 10) identified from prey proteins. This result provides evidence that SAAP originate from the proteins to which we have assigned them. Despite the significant difference between the primary tissues and HEK cells, the corresponding ratios between SAAP and BP correlated significantly, Supplemental Fig. 5c. This correlation is naturally weakened by factors such as HEK cell specific rates of SAAP production and degradation rates differences. The IP-MS spectra contained numerous fragments supporting the SAAP sequences with intensities corresponding to Prosit predictions, Supplemental Fig. 5d.

As a final validation of SAAP sequences, we used PepQuery^30^ to run a peptide-centric search of the substituted peptides against the dataset spectra. On average, 78% of substituted peptides were confidently matches to spectra, a rate similar to that of the base peptide positive controls. Notably, a negative control set of reversed substituted peptide sequences matched to spectra at an average rate of only 1%, Supplemental Fig. 6a.

### Quantification of amino acid substitutions

To provide context for the prevalence and possible functions of SAAP, we investigated the abundance of SAAP relative to their corresponding encoded base peptides (BP). The abundance of a peptide depends both on the synthesis rates and the degradation rates of its parent proteins. At steady state, the ratio between the peptides with and without substitutions, i.e. SAAP/BP, which we term RAAS, equals the ratio of corresponding synthesis and degradation rates of the alternatively translated and encoded proteoforms, Fig. 2a. Most SAAP are expected to be synthesized at a low rate and degraded at a high rate, resulting in low SAAP abundance and low RAAS. However, even substitutions incorporated at low rates may accumulate to high level if they stabilize the resulting proteoforms^31^.

While median RAAS estimates are consistent with previous reports (reviewed by ref.^12^), we observed that about 10% of SAAP have substitution ratios exceeding 1, Fig. 2b. Surprisingly, our estimates suggest that the alternatively translated proteoforms are the most abundant protein products for 360 proteins spanning diverse biological functions. This finding is highly unexpected and motivated us to further evaluate it based on multiple lines of evidence. First, a detailed inspection of the underlying RNA sequence data and mass spectra provide strong support for the absence of alternate alleles and the confidence in the assigned amino acid sequence, Supplemental Fig. 1 and 2. Accordingly, we find that the identification confidence to be the same for SAAPs having high and low RAAS, Extended Data Fig. 2a-c. Second, we tested if differences in peptide ionization can explain high RAAS and found that SAAP have about the same ionization efficiency as their corresponding BP according to the model proposed by Liigand, *et al.*^33^. Differences in a single amino acid residue have modest effects on ionization, and even the largest deviations are too small to significantly affect RAAS estimates, Extended Data Fig. 2d-f. We similarly computed peptide detectability using the DeepMSPeptide method described by Serrano, *et al.*^34^. As shown in Extended Data Fig. 2g-i, peptide detectability is not significantly different between substituted and base peptides. There is no significant correlation between RAAS and peptide detectability (*r* = −0.0004, *p* = 0.97). Third, we confirmed strong agreement in RAAS estimates for the same substitution quantified from different peptides originating from independent digestions with different proteases, Fig. 2c, Extended Data Fig. 2j. Fourth, we ensured that high RAAS are not an artifact of having highly abundant BPs with missed cleavages, Extended Data Fig. 2k. We also confirmed strong consistency between RAAS values for peptides confidently identified by Pep-Query^30^ and those not matched, Supplemental Fig. 6b. Thus, five lines of evidence support the accuracy of our RAAS estimates.

As a sixth validation of RAAS estimates, we tested whether they are consistent with the abundance of peptides shared between canonical and substituted proteoforms, as estimated from their precursor intensities. The abundance of the shared peptides (other peptides from the same protein) should reflect the cumulative abundance of both encoded (P1) and alternatively translated (P2) proteoforms, Fig. 2a, which is approximated by the most abundant proteoform, especially for very high and low RAAS. This expectation is strongly supported by the data, as illustrated by an example of a highly abundant substitution (RAAS*>*6) identified with high-confidence (positional probability*>*0.9) in the fructose-1,6-bisphosphatase 1 protein, Fig. 2d. Consistent with the high RAAS, the abundance of the shared peptides is closer to the SAAP abundance than to the BP abundance. To generalize this analysis to the entire dataset, we computed the ratios between the abundances of shared peptides and SAAP or BP, Fig. 2e. The results indicate that for peptides with low RAAS, the BP peptide abundance is similar to the shared peptide abundance, while the SAAP abundance is orders of magnitude lower, Fig. 2e (bottom panel). As RAAS increases, the abundance of the SAAP becomes less distinct from the abundance of the shared peptides, while the BP abundance increasingly deviates from the abundance of shared peptides, Fig. 2e (top panel). For RAAS*>*1, substituted and shared peptides have similar abundance, Fig. 2d,e. Together, these trends bolster the conclusion that P1 proteoforms are dominant at low RAAS while P2 proteoforms are dominant at high RAAS.

Our final test for the reliability of high RAAS evaluated their consistency with the correlation patterns of BP or SAAP abundance to shared peptide abundance across patients, Extended Data Fig. 2l. These six complementary analyses indicate that the quantification derived from many peptides of different types (BP, shared peptides, and SAAP) at both MS1 and MS2 level all dovetail together and support that SAAP with RAAS*>*0 represent peptides from the most abundant proteoforms. When additional stringent filters are applied to remove potential biases associated with missed cleavages and substitution position, the majority of identified substitutions remain valid, Supplemental Fig. 7. Therefore, thousands of proteins have abundant proteoforms with amino acid substitutions, including hundreds with confidently localized substitutions. For some proteins, these are the most abundant proteoforms.

Having established confidence in the estimates of substitution ratios, we explore them across the different proteins, patients and cancer types. These ratios span 10^8^ range at the level of individual substitutions (Fig. 2b and Extended Data Fig. 3a,b) which shrinks to 100-fold range across patients (the median RAAS per patient), Extended Data Fig. 3c, and to 4-fold across cancer types, Extended Data Fig. 3d, Supplemental Data Tables 3,4. While most observed SAAP are specific to a dataset, there is notably higher overlap between the two lung cancer datasets and a cluster of shared SAAP that are also commonly found across the majority of patients in a dataset, Extended Data Fig. 3e,f. RAAS estimates based on peptides found in all datasets, which are not affected by differences in peptide detection, show significant (*p <* 10*^−^*^12^) variation in RAAS across cancer types, Extended Data Fig. 3g,h. Further, ratios of RAAS for SAAP intersected between pairs of patients show significant (*p <* 10*^−^*^20^) differences across datasets, with medians of relative RAAS varying over 2-fold from the lowest to the highest, Fig. 2f,g. These estimates suggest that there is biologically-driven RAAS variability across proteins, cancer types, and patients.

The proteins with high substitution ratios span multiple functional groups (Fig. 2h), including signal transduction, protein degradation and transcriptional regulation. Estimation of protein copy numbers by the histone ruler method^32^ suggests that highly abundant alternatively translated proteoforms are present at hundreds to tens of thousands of copies per cell, Fig. 2i.

### RAAS depends on codon frequency, tRNA pairing and RNA modifications

Confident in our quantification of SAAP, we explored the dependence of RAAS on the number of nucleotide mismatches corresponding to a mRNA-tRNA pairing that can translate the detected SAAP. This analysis indicated a strong dependence between RAAS and the minimum number of codon mismatches required for the corresponding substitution, Fig. 3a, an effect consistently observed across the datasets, Extended Data Fig. 3i. Just as 3 base pair mismatches are less likely to occur, so too substitutions that would require complete codon mismatch have lower RAAS. We also observed that for some substitutions, such as *T* → *V* in cancer, there is significantly higher abundance of the tRNA ligase loading the incorporated amino acid relative to the tRNA ligase loading the encoded amino acid, Extended Data Fig. 3j.

Exploring the dependence of substitution ratios on codons, we found that they are inversely proportional to relative codon frequency (Fig. 3b and Extended Data Fig. 4) and to the codon stability coefficient (Extended Data Fig. 3k), which is an empirical measure of codon usage^35^. These observations are consistent with a previously proposed model^36,37^ that less frequent codons are associated with a smaller tRNA pool, which may result in increased rate of substitution by the ribosome for a tRNA that is more readily available, increasing alternate decoding at these sites.

Furthermore, we explored RAAS dependence on RNA modifications detected by direct RNA sequencing using nanopores. Using data by McCormick *et al*^38^, we found that uracil modifications overlap significantly (*p <* 10*^−^*^10^) with substitution sites (Supplemental Fig. 8a) and the modification fraction correlates significantly to the substitution ratios, Supplemental Fig. 8c,d. Together, these data suggest that multiple RNA related mechanisms influence the rate of alternate decoding, and thus contribute to high substitution ratios.

### Increased protein stability contributes to substitution abundance

While variation in the rate of alternate translation clearly contributes to RAAS, protein stability may contribute as well, Fig. 2a). Fortunately, this contribution can be directly quantified using metabolic pulse with stable isotope labeled amino acids. Thus, we analyzed data from such experiments with primary human liver-derived cells^39^. Applying the analysis pipeline from Fig. 1a, we identified over 10,000 SAAP with a RAAS distribution similar to the one observed from the TMT and label-free data, Extended Data Fig. 3l. These similarities extend to detecting SAAP with identical sequences and substitutions as those detected in the other datasets, Supplemental Data Table 5. SAAP detected in hepatocytes have 5-fold higher overlap with SAAPs from the label-free healthy liver samples compared to other tissues (Supplemental Fig. 9a), supporting prevalence of alternate translations in tissue-specific proteins. We validated these findings with an independent database search using the MSFragger pipeline^40^, with deep-learning rescoring enabled^41^, Supplemental Fig. 9b. These results generalize our observations across another type of MS data acquisition that offers further constraints (from stable isotope incorporation) on sequence identification.

Our analysis of the metabolic pulse data allows direct evaluation of the dependence between protein degradation rates and RAAS. Many substituted peptides have lower degradation rates than their non-substituted counterparts, Fig. 3c. This global trend is reflected in a strong inverse correlation (*p <* 10*^−^*^10^) between RAAS and degradation rates in hepatocytes, Fig. 3d. Similar inverse correlations are observed with primary B cells and natural killer cells, Extended Data Fig. 3m,n.

To further test the impact of substitutions on protein stability, we extended the analysis to shared peptides, analogous to the abundance analysis in Fig. 2d,e. To exemplify this with a specific protein consider prolyl endopeptidase: it has a high ratio substitution and the SAAP has stability similar to the shared peptides, while the BP has a high degradation rate, Fig. 3e. Global analysis of the ratios between the degradation rates of SAAP to shared peptides and between BP to shared peptides degradation rates support the generalization of this finding, Fig. 3f. On average shared peptide degradation rates for proteins with lowly abundant substituted proteoforms are more similar to the degradation rate of the encoded peptide (log ratio∼0), while the substituted peptide degradation rate is higher (positive log ratio). For proteins with highly abundant substituted proteoforms, the shared peptides degradation rates are more similar to the degradation rate for the substituted proteoform (log ratio∼0), while the encoded peptide degradation rate is higher (positive log ratio). The degradation rate ratio difference between the two RAAS bins is statistically significant for both substituted (p=0.00015) and base peptides (p=0.0098).

This degradation analysis is highly complementary to the abundance analysis and bolsters our results. The effect size for the degradation analysis is smaller since protein stability variation is only one of the factors contributing to RAAS. Second, the dataset used to compute turnover has shallower coverage. For these two reasons, the the signal to noise in the degradation rates is smaller compared to the abundance measures. Nonetheless, the results strongly support that for high RAAS, the turnover of the shared peptides is closer to that of the SAAP (not the BP), consistent with the substituted protein being the main proteoform.

These results indicate that many substituted proteoforms with high RAAS are stable, i.e., have low protein degradation rates. This is consistent with the expectation that most substitutions likely destabilize proteins and result in low abundance SAAP, below our detection limit. Yet, the SAAP that are detected, particularly those with high RAAS, correspond to substitutions that increase the lifetimes of their proteoforms in living cells, Fig. 3d.

### Substitution ratios depend on amino acid and tissue types

Next, we examined how RAAS depends on the substitution type defined by the specific combination of encoded and incorporated amino acids. We find that the median substitution ratios for different substitution types vary from 10*^−^*^4^ to over 1, Fig. 4a,b and Extended Data Fig. 5a. Thus substitution type can explain much of the observed variation in substitution ratios. Still, ratios vary within a substitution type as well; this variation is relatively small for some types, such as *H* → *Q*, and larger for others, such as *E* → *D*, Fig. 4a. The majority of substitution types are represented by SAAP that are over 100-fold less abundant than the corresponding encoded BP, Fig. 4a and Extended Data Fig. 5b. Yet, some substitution types, such as *S* → *G*, have median RAAS*>*1, Fig. 4a,b. Most substitution types with RAAS*>*1 are observed relatively infrequently in the data, and their average properties therefore are more influenced by individual SAAP, Fig. 4a and Extended Data Fig. 5a. Interestingly, the substitution ratios of substitution types cluster by the chemical properties of the encoded amino acid, Fig. 4b,c and Extended Data Fig. 5c. Specifically, substitutions of polar amino acids have consistently higher RAAS than substitutions of other amino acid types. This effect is strongest for polar → special substitutions. In contrast, substitutions of charged or hydrophobic amino acids result in SAAPs with lower ratios, Fig. 4b,c and Extended Data Fig. 5c.

The pairwise correlations of substitution-type median substitution ratios across datasets indicate significant similarities, with no obvious global distinction between the label-free (Healthy) data and the TMT-labeled CPTAC data, Fig. 4d. Variance in substitution ratios for individual substitution types across datasets is highly dependent on substitution type. We find that some substitution types, such as *E* → *N* have significant agreement in RAAS across datasets, Extended Data Fig. 5d, while others, such as *P* → *S* show clear dataset dependence, Extended Data Fig. 5e. Similarities in substitution ratios of substitution types across datasets is majorly driven by similarities in RAAS for SAAP with the same encoded amino acid, rather than RAAS consistency for incorporated amino acids, Fig. 4e and Extended Data Fig. 5f,g. These findings support significant dependence of RAAS on the substitution type, and we sought to quantify it using ANOVA. The results (Fig. 4f) indicate that substitution type, especially the encoded amino acid, explains the most variance in RAAS. Tissue type is also significantly associated with substitution ratios. Fold change analysis of RAAS for substitution types in a specific tissue (using both cancer and healthy data) relative to all other tissues indicates that some substitutions, such as *G* → *S* in pancreas, have significantly (*p <* 10*^−^*^15^) tissue-specific RAAS distributions, Extended Data Fig. 5h,i.

### Linking substitutions to protein function and structure

Having established the associations of substitution ratios with substitution type and tissue specificity, we next explored potential structural and functional consequences of alternate translation events, starting with differences between cancer and adjacent non-cancer tissue. While we did not have the power to detect global trends in RAAS with tumor status (Extended Data Fig. 6a,b), there are clear outlier SAAP for which substitution ratios in some tumor types are significantly higher than RAAS in adjacent non-cancer from the same patient. One striking example is a substitution with strong fragment ion evidence in lamin isoform A protein, for which RAAS in tumor samples was significantly higher than in surrounding control tissues in 3 cancer types, Fig. 5a. Another example is the *N* → *G* substitution in the serine/threonine-protein phosphatase PP1-beta catalytic subunit. For lung cancer patients, this substitution has consistently higher RAAS in tumor samples than in the matched controls, Extended Data Fig. 6c. Furthermore, specific substitution types, such as *H* → *N* and *T* → *Q* exhibit significantly different RAAS in tumor samples compared to matched surrounding tissues, Fig. 5b. Together, these results demonstrate cancer specific differences in the abundance of protein products of alternate RNA decoding.

To identify biological functions that are most affected by alternate translation, we next identified Gene Ontology (GO) protein sets enriched with highly abundant alternatively translated proteoforms, Fig. 5c and Extended Data Fig. 7. These significant GO groups suggest a broad impact of substitutions on the proteome, including proteins functioning in gene expression, cellular organization, and signaling. As suggested by observations that alternatively translated proteasome subunits are highly abundant, Fig. 2h, protein catabolic processes are significantly enriched with high RAAS proteoforms, Fig. 5c, Extended Data Fig. 7. Interestingly, signaling proteins are also enriched for high RAAS, indicating potential downstream consequences, Fig. 5c and Extended Data Fig. 7. For example, an alternatively translated isoform of EZR is found in all datasets, which may affect cellular adhesion and migration processes.

To explore the context of substitutions, we investigated sequence patterns associated with alternate translation. Sequence logos and amino acid enrichment around substitution sites and along tryptic peptides revealed four enrichment patterns common to different substitution types, Extended Data Fig. 8a-d. We observed that many highly abundant SAAP have a *Q* → *G* substitution.

While the mass shift for this substitution is also consistent with A cleavage from the peptide N-terminus, the fragment ion intensities are better explained by *Q* → *G*, Supplemental Fig. 4b-f. Through sequence motif analysis, we identified 768 SAAPs significantly enriched for a “KRAQ” motif, in which a K, R, A, and G are enriched immediately before a *Q* → *G* or *Q* → *A* substitution, Extended Data Fig. 8d. This sequence motif has substitution ratios of about 0.1 across all datasets (Fig. 4c bottom panel), significantly (*p <* 10*^−^*^10^) above the median RAAS.

Furthermore, we found that substitutions surrounded by CC, MM, or WW tend to have high RAAS and all show enrichment for specific substitution types, Extended Data Fig. 8d. For example, CCxCC corresponds to high probability of Q or N to be substituted by A or G. Additional motifs with high RAAS substitutions, include an enrichment of M in +/-1 and +/-2 positions relative to the substitution site, Extended Data Fig. 8d. This motif is generally found in the context of substitutions of T and S. A spatial proximity between reversibly sulfoxidized M and S/T kinase targets has been previously noted^42^, but a biological function or effects on protein stability of these loci have not been described. Another similarly interesting motif identified is W in +/- 1/2 position relative to the substitution, Extended Data Fig. 8d. W is both the rarest and metabolically most costly amino acid^43^, and tryptophan codons have been observed to cause ribosome stalling and frame-shifting translation errors (“W bumps”) in melanoma^44^. Here, we find W in the context of substitutions of branched chain amino acids, which are themselves known to be dysregulated in starvation and diabetes^45^, and particularly in PDAC^46,47^. While these results suggest the existence of amino acid sequence patterns that are significant targets of alternate translation, we cannot confidently rule out the possibility of multiple modifications or alternate events resulting in the relevant mass shifts for these putative motifs.

These observations led us to explore more broadly the spatial clustering of substitutions, both within structural domains and within regions having high density of substitutions in the primary amino acid sequence (1D) or in the 3D structure of proteins. Analysis of Pfam domains confirmed enrichment of abundant substitutions in several structural domains, including those related to significant functional protein sets, such as the proteasome subunit domain, Ras family domain (signaling), and RNA recognition motif, Extended Data Fig. 9a. Many substitutions occur close to each other, with over 1,200 base peptides having multiple substituted peptides, Extended Data Fig. 9b. An example of such clustered substitutions is shown in Fig. 5d with the significantly enriched thioredoxin domain, Extended Data Fig. 9a. The ratios of these substitutions increase 100-fold over 10 amino acids, Fig. 5d. Further examples of clusters of substitutions with both high and low RAAS occur in the actin domain of ACTG2 (Fig. 5d), as well as in MZB1 and the glycolytic domain of ALDOB, Extended Data Fig. 9c,d. We were also able to identify structural domains with clusters of alternate translation in 3D. Notably, the significant Ras family domain (Extended Data Fig. 9a) has three substitution sites with high RAAS that cluster together in the 3D structure of the RAP1A protein, Fig. 5e. Other interesting domains with clusters of alternate translation include the ribosomal and proteasomal protein complexes, Extended Data Fig. 9e,f.

### Predictors of substitution ratios

To place alternate translation in broader biological context, we examined protein attributes correlated with median RAAS. We find that RAAS is positively correlated to protein length, Fig. 5f, and negatively correlated to protein half-life, Fig. 5g, features which are known to negatively correlate to one another by thermodynamic principles^48^. This suggests that shorter, more stable proteins, are less likely to have alternatively translated proteoforms that are more stable than their encoded form, with the caveat that these features and RAAS are also correlated to protein abundance, Extended Data Fig. 10a. Additionally, we observe that substitution sites with high RAAS are positively correlated with protein regions that are more highly disordered and inversely correlated with more lowly conserved regions, Fig. 5h and Extended Data Fig. 10b,c, in line with the expected inverse correlation between conservation and disorder. This intuitively suggests that more highly conserved and structured protein regions are less likely to have alternatively translated proteoforms that are more stable than their canonical counterparts. These features, along with substitution site position and binding site information were used to successfully predict RAAS distributions using a random forest model, Fig. 5i. While these trends are broadly informative their interpretation has a caveat: Low ratio substitutions on lowly abundant proteins are less likely to be detected, leading to overestimation of the *median* RAAS for lowly abundant proteins, which may induce indirect correlations of features associated with protein abundance.

The strong association of RAAS values with protein sequence conservation led us to explore the association with allele frequency variation in the human population using the gnomAD database^49^ (see Methods), Supplemental Data Tables 7 and 9. We estimated the population frequency of missense variants in alternatively translated codons. The results for all possible allele variants within alternatively translated codons (Fig. 5j) and for the subset of nucleotides that may lead to the incorporated amino acid (Extended Data Fig. 10d) indicate that most alternate translation sites have no observed allele variants in gnomAD. This result provides further evidence that the substitutions identified in our data do not arise from genetic polymorphisms translated according to the genetic code. Furthermore, we found that codons corresponding to high RAAS substitutions have more allele variation than expected, suggesting that they are less constrained than codons corresponding to low RAAS substitutions, Fig. 5k. This trend is even stronger when considering only the subset of alleles that may encode the incorporated amino acid, Extended Data Fig. 10e. This result reinforces the association with protein conservation, namely that alternate translation occurs with higher frequency at sites that are more tolerant to sequence variation.

### Conservation across species

Lastly, we assessed conservation of alternate translation across species. We ran our pipeline from Fig. 1a on label free proteomics data from 3 mouse tissues^14^ and identified 1,102 substitutions events representing 397 unique SAAPs, 55 of which are identical with human SAAPs analyzed above, Extended Data Fig. 10f. This intersection demonstrates conservation, though it likely underestimates the degree of conservation since (1) we did not account for protein sequence homology differences and (2) technical factors affect peptide detectability. To more quantitatively assess the conservation of alternate translation, we further analyzed RAAS values. Their significantly high correlation (*p <* 10*^−^*^8^) across human and mouse strongly support species conservation, Fig. 5l. The total variance in substitution ratios of these shared SAAP are most significantly driven by substitution type, followed by tissue type, Fig. 5m.

## Discussion

Our results demonstrate that alternate translation produces thousands of stable and abundant proteins in human and mouse tissues. These proteoforms differ in sequence, similar to proteoforms resulting from missense mutations. However, there are important differences. First, mutations introduce about 44 missense substitutions per cancer^50^ while alternate translation introduces thousands of sequence changes. Second, mutations affect all protein copies templated by the mutated allele while alternative decoding affects only a fraction, though the fraction can be large and amount to O(10^4^) copies per cell, Fig. 2h. Furthermore, some substitutions are significantly more abundant in tumors (Fig. 4a), suggesting a disease association. Another potential link to disease are the highly abundant substitutions in numerous proteins associated with neurodegenerative diseases, such as TDP43, FUS and VCP. These substitutions are identified across all datasets and are reported in Supplemental Data Table 6.

Multiple mechanisms likely contribute to the amino acid substitutions that we quantified. Many substitutions, especially those with low ratios, likely reflect limited fidelity of aminoacyl-tRNAs recognizing their cognate codons. This mechanism is supported by higher ratios for aminoacyl-tRNAs pairing based on fewer nucleotide mismatches (Fig. 3a and Extended Data Fig. 3i) and for rare codons, Fig. 3b, Extended Data Fig. 3k, and Extended Data Fig. 4. High ratio substitutions likely involve sense codon recoding that may involve tRNA and mRNA modifications. Indeed, mRNA pseudouridylation recodes stop codons^17,18^ and cytidine acetylation affects translational fidelity^51^. We found that U modifications overlap significantly (*p <* 10*^−^*^10^) with the amino acid substitutions reported here, Supplemental Fig. 8. This is consistent with its potential to recode sense codons previously observed with exogenous synthetic constructs^52^. Such recoding might also be regulated by UTR sequences altering codon pairing analogous to the way the Selenocysteine Insertion Sequence alters the reading of the UGA stop codon into incorporating selenocysteine^53^, though our data do not provide direct evidence for such UTR sequences. Similarly, modifications of transfer RNA, ribosomes^54–56^, RNA binding proteins and structures delaying elongation may contribute to sense codon recoding^57^. A major mechanism determining the abundance of the substituted proteins is their stability, which we directly estimated based on protein degradation rates, Fig. 2.

Our results reveal only a subset of alternatively translated proteins because (1) we tested a limited sequence space (a single amino acid deviation from genetic code prediction), (2) we analyzed only a subset of all peptide ions, and (3) limitations exist in interpreting the mass spectra of analyzed ions^58^. Indeed, even the deepest MS datasets cannot achieve full protein sequence coverage^59^. In the data-dependent-acquisition datasets analyzed here, the peptide ions were selected for fragmentation in order of their abundance, and therefore less abundant peptides are less likely to be fragmented and identified. As a result, lowly abundant substituted peptides are less likely to be identified and their ratios are more likely to be missing from our data; thus, lower ratios are under-represented in our RAAS distributions. Even more limiting, only a minority of all fragmented peptide ions are assigned to confident sequence, which is a general challenge in MS proteomics^8,9,60^. This sequence assignment remains limiting when all detectable ions are sampled by data-independent-acquisition methods, as current methods can assign confident sequence to less than 25% of the peptide ions^61^.

The sequence assignment challenge is heightened by our priority on minimization of false positives, which likely increased false negatives. For example, we excluded 2,734 SAAP whose sequence might correspond to alternative reading frames and DNA sequences annotated as noncoding. Yet this hypothetical sequence correspondence is unlikely to represent a templating relationship. We also excluded thousands of SAAPs that did not pass our very stringent filters (Fig. 1) even though they are likely correctly identified. We chose these conservative and rigorous filters to increase confidence in the remarkably large number of abundant SAAPs. The mass spectra of some SAAPs with high ratios contain fragment ions corresponding to fragmentation of each peptide bond and are thus consistent with only one amino acid sequence, Supplemental Fig. 1 and 2. However, some mass spectra are less complete and provide lower confidence. The substitutions of such sites are further supported by fragmentation patterns (Supplemental Fig. 4) and their significant associations with mRNA modifications (Supplemental Fig. 8), protein disorder, conservation, and allele polymorphisms, Fig. 5. Future research is needed to increase the number of detected fragments^62^, to balance the trade-offs between false positives and negatives, improve estimated of false discovery rates of SAAPs, and expand the set of confident SAAP whose biological significance can be interpreted.

Due to the above-mentioned limitations of our analysis, it is possible that alternate hypotheses such as multiple modifications on the peptide can explain the mass shifts observed for some of our substituted peptides. Since it is practically infeasible to disprove every alternative hypothesis, we have focused instead on providing confidence metrics based on the localization probability of the identified substitutions. We demonstrate that even with the most stringent filtering of substitutions based on confidence, we still identify 590 unique substituted proteoforms with RAAS exceeding 0.1 that are likely functionally relevant. These high-confidence and highly abundant substitutions are provided in Supplemental Table 8.

Sites on lowly abundant proteins with low RAAS are less likely to be detected. This may results in non-ignorable missing data for some analysis, such as establishing the biological association between median RAAS per protein and protein abundances. To mitigate these confounding effects, we focused our conclusions on ratios between quantified substitution sites to establish ratio variation across patients and cancer types, Fig. 2e,f and Fig. 4a,b.

Protein products of alternate translation are challenging to detect since most interpretation of MS data is based on database-searching that tests hypotheses about the detection of protein products whose sequence matches nucleic acid sequences; hypotheses about the presence of protein products with sequences not predicated by the genetic code are not commonly tested. Furthermore, if such proteins are identified, they need to be further evaluated by a battery of additional tests, and downstream analysis as reported here. For example, previous reports of substitutions^14,58^ did not test if they have genetic origin and did not focus on their validation, abundance and stability. We expect that advances in *de novo* protein sequencing and increased protein sequence coverage will identify many more substitutions. The abundance of many substitutions reported here is high enough (Fig. 2) to quantify them in single cells^63,64^, especially if their analysis is prioritized^65^.

Our results provide direct evidence for the abundance and stability of proteins with amino acid substitutions (Fig. 2) and associative evidence for their functional roles, Fig. 4 and Fig. 5. To avoid assuming functions or mechanisms, we chose the neutral phenomenological term “alternate translation”. While some SAAPs may merely reflect limits of translational fidelity and proteostasis, others may have evolved biological functions as previously suggested^15,16^, consistent with regulated sense codon recoding discussed above. The high abundance, conservation across species, and associations of some SAAPs with cancer and protein domains (Fig. 4) imply functional significance, and this possibility needs to be explored directly by future research.

The conservation of SAAPs between human and mouse (Fig. 4) contrasts with limited conversation between mammals and unicellular microorganisms, including yeast and bacteria. Previous analysis of substitutions in yeast and bacteria did not discover highly abundant substitutions^13^, and we confirmed this result.

## Methods

### Sample-specific protein databases

RNA-sequencing data was downloaded from NIH Genomic Data Commons (CPTAC data, portal.gdc.cancer.gov) or from dataset repository (label-free data, E-MTAB-2836). Reads were sorted by name and paired ends were written to two .fastq files (CPTAC only) with samtools v1.10^66^. Adapters were trimmed, reads were filtered for phred quality ≥ *Q*28 and QC data was printed with fastp v0.23.4^67^. The remaining reads were aligned to GRCh38 reference genome with hisat2 v2.2.1^68^ with –dta flag for downstream trancriptome assembly processing. The alignment was converted from .sam to .bam, sorted by position and indexed with samtools v1.10^66^. samtools depth v1.10^66^ was also used to get the read depth at each base. Aligned reads were processed into a custom protein database with a 2-pronged approach: De-novo transcriptome assembly and single nucleotide variant (SNV)/insertion-deletion (indel) calling. First, the sorted and indexed hisat2 alignment was used to assemble a de novo transcriptome with stringtie v2.2.1^69^, which captures alternative splice variants. A transcript was called at a given locus if its abundance was at least 0.1% of the most abundant transcript at that locus, and if there was at least 1 independent read for that transcript. The assembly was compared to the reference genome using gffcompare v0.11.5^70^ and the annotated result was filtered for coding regions (CDS) with gffread v0.12.7^70^ and converted into .bed format using a Galaxy-sourced python script (gffcompare to bed.py)^27^. A second Galaxysourced script (translate bed.py) was used to translate the CDS regions of the annotated de novo assembled transcriptome into protein sequences (based on the genetic code) and generate a fasta database of the translated sequences. Second, SNV and indel variants were called using freebayes v1.3.4^71^ with the sorted hisat2 alignment as input, and default parameters. customProDB v1.41.0^72^ was then run to translate the SNV and indel calls into protein fasta databases. Fasta databases from the two variant calling processes were merged into a single protein database for each sample using another Galaxy-sourced python script (fasta merge files and filter unique sequences.py)^27^. Sample-specific protein databases for all samples in a TMT experiment (CPTAC data) or all samples of a given tissue type (label-free data) were further merged. This pipeline was implemented on the Linux command line using the Discovery high performance computing cluster (MGHPCC, Holyoke, MA).

### Blast

All proteins in the custom protein databases were blasted against the RefSeq human proteome using the blastp command from ncbi-blast+ command line tool at: ftp.ncbi.nlm.nih.gov/blast/executables/blast+/2.16.0/). The top matches for each protein were used to identify the most likely source of ambiguous proteoform translations.

### Proteomics data processing

LC-MS proteomics data was downloaded from NIH Proteomic Data Commons (CPTAC data, proteomic.datacommons.cancer.gov/pdc/) or from dataset repository (label-free human data, PXD010154, and label-free mouse data, PXD030983), in the form of .raw files. For the discovery search, raw LC-MS data for each TMT set (or tissue for label-free data) was searched against the corresponding custom protein database using MaxQuant v1.6.17.0^6^ with dependent peptide (DP) search option set to True and dependent peptide FDR set to 1%. For the validation search, the SAAP sequences were appended to the sample-specific FASTA file, the DP search option was set to False, and match between runs was enabled. PSM, peptide and protein FDR were all set to 1%. To run DP search with TMT-labeled data, we set LCMS run type to “Standard” and included TMT labels as a fixed modification at the N-terminus and lysine residues of peptides. To search label-free data, the TMT modification was omitted. Cysteine carbamidomethylation was included as a fixed modification, and methionine oxidation and N-terminal acetylation were added as variable modifications. Other parameters were set as defined by the original publications and were consistent across CPTAC datasets. See project Github for a sample configuration file. Searches were implemented on the Linux command line using the Discovery high performance computing cluster (MGHPCC, Holyoke, MA).

### Candidate SAAP and PTM identification

DP search results were mined for peptides exhibiting a mass shift that corresponds to either a known PTM (unimod.org) or to a potential AAS using custom python scripts including functions adapted from ref.^13^. Only a single modification per peptide was considered. The mass error threshold between the observed mass shift and the mass shift of a given modification was required to be below 5 ppm. For PTMs, we required that an amino acid known to be modified by that PTM was present in the peptide sequence. For AAS, we further required that that amino acid was assigned a positional probability of modification by MaxQuant. Peptides that could be explained by both an AAS and a PTM were not considered as candidate SAAP, e.g., we filtered out *Q* → *E* and *N* → *D* as such changes may arise as part of deamidation. Base peptides were allowed to have more than one associated modified peptide. The few identified peptides with substitutions of K and R are filtered out due to the potential impact of miscleavage on abundance estimation. Peptide sequences with potential AAS were compared against a six-frame translation of all regions of the human genome and any matches were discarded as potential products of non-canonical translation.

### Mass shift localization

When the MS2 spectra of SAAPs contain fragments generated from cleavages of the precursor ions just before and just after the substituted amino acid, the mass shift can be confidently assigned to a single amino acid residue. The confidence of such localization is reflected in the positional probability estimated by MaxQuant, Supplemental Fig. 4a. When the positional probability is close to 1, the measured mass shift can be confidently assigned to single residue, a substituted amino acid or another modification of the residue. However, MS2 spectra often do not contain a full set of fragments, especially from lowly abundant peptides, sequences that fragment poorly or small fragment ions (e.g., b1) below the low limit of MS2 spectra.

In such cases, we used fragmentation patterns to determine whether an observed mass spectrum with uncertain positional probability aligns better the proposed SAAP sequence or with an alternative hypothesis that can explain the observed mass shift relative to the BP. Specifically, we predicted the fragment ion spectra for the SAAP and the peptide representing the alternate hypothesis using Prosit^73^ as implemented in Koina^74^ and shown in Supplemental Fig. 4b-f. For TMT-labeled peptides the `Prosit 2020 intensity TMT` model^75^ was used with HCD fragmentation, and for label-free peptides the `Prosit 2020 intensity HCD`^73^ model was used. Predicted spectra for the SAAP and the alternate hypothesis peptides were compared to the empirical spectra with a cosine similarity score computed with the `CosineGreedy` function from the matchms.similarity python package^76^, 4. Koina was also used to predict spectra for high-confidence SAAPs and their corresponding BPs as shown in Supplemental Fig. 1 and 2.

### Rescoring SAAP confidence

To validate if the discovered SAAP represent genuine peptides that can not be explained by more confident canonical PSMs, closed validation search outputs from MaxQuant v2.4.3 for the entire CCRCC cohort were rescored using the Prosit model and Percolator within the Oktoberfest pipeline^8^. Unfiltered database search results performed at 100% FDR including decoy PSMs as well as raw files were used as input. Normalized collision energy (NCE) calibration of the model was performed in the range 19-50. Peptide intensity prediction was performed with the model “Prosit 2020 intensity HCD” and indexed retention times were predicted with “Prosit 2019 irt”. Matching of predicted intensities was performed using previously described similarity measures such as spectral angle and Pearson correlation. False discovery rate was estimated using the SVM Percolator^77^ with its standard settings.

### Sequence validation with PepQuery

To validate the novel SAAP sequences, we made use of the peptide-centric PepQuery search^30^. The standalone version of PepQuery (v2.0.2) was installed on the Discovery Cluster. Raw proteomics data files for LUAD and LSCC were converted to .mgf files using msconvert**^ProteoWizard^**. Before searching the sequences against the spectra, the mfg files were used to create an index using the PepQuery index function. Three searches were done for each of the two lung cancer datasets: SAAP sequences, BP sequences,reversed SAAP sequences. A text file with the peptide sequences and the directory location of the pre-created index were used as input (-i and -ms parameters, respectively) to the pepquery-2.0.2.jar function call. For SAAP and reverse SAAP sequences, which served as a negative control, a novel peptide validation search type was conducted (-s 1). For the positive controls of encoded (BP) sequences, a known peptide validation search type was conducted (-s 2). Other parameters were the same for all searches. The human uniprot fasta database was input as the reference. Cysteine carbamidomethylation and TMT labeling of lysine and arginine were set as fixed modifications; Methionine oxidation and N-terminal acetylation were set as variable modification. An example PepQuery function called is provided below.

`java -jar pepquery-2.0.2.jar -i LSCC PepQuery seqs.txt`

`-t peptide -s 1 -ms /scratch/tsour.s/Sat LSCC/Proteomics data/index`

`-db uniprotkb human.fasta -fixMod 1,11,12 -varMod 2,5`

`-o LSCC saap -plot`

Peptides were considered to be confidently matched to a spectra by PepQuery if the relevant ”confident” filter in the psm rank.txt output had a value of ”Yes”.

### Quantifying SAAP, BP, and RAAS

For CPTAC datasets, peptide abundance was computed at 2 levels: precursor and reporter ion. The abundance of precursor ions corresponds to the total intensity of a peptide across all samples in a given mutiplexed TMT-labeled experiment (experiment level). This value is reporter by MaxQuant in the “Intensity” column of the evidence.txt output file. Reporter ion abundance corresponds to the intensity of a peptide in an individual patient sample (sample-level). This value is computed by distributing the precursor ion intensity by the fractional contribution of each reporter ion in the experiment to the total reporter ion intensity for that peptide. This method was used to calculate abundance of all peptides reported here, including SAAP, BP, and other peptides. For the label-free dataset, only precursor-level peptide abundances were computed. If the peptide is quantified in multiple charge states and across different off-line fractions, the intensities of all instances were summed to estimate the overall abundance. For peptides other than SAAP and BP, modified versions of the peptide were included in the sum. Ratios of amino acid substitution (RAAS) were computed by dividing the SAAP abundance by the BP abundance and are generally reported on a log_10_ scale in figures and supplemental tables. If log10 is not specified, RAAS is reported on a linear scale in that instance (occurs throughout text). If the substitution impacts the cleavage probability, the distribution of the peptide across fully cleaved and miscleaved forms may be altered and impact RAAS estimation. To limit the impact of such artifacts, we removed from our analysis all base peptides that were also identified as part of longer miscleaved peptides, Extended Data Fig. 1h(i). Precursor ion, experiment-level SAAP quantification data can be found in Extended Data Table 3. Reporter ion, sample-level SAAP quantification data can be found in Extended Data Table 4.

### Analysis of IP-MS samples

IP-MS experiments for 25 proteins with SAAPs detected from human tissues were downloaded from the bioplex project^29^. Raw files were searched against a human protein sequence fasta downloaded from Uniprot with all SAAPs identified from human tissues listed in Supplemental Table 2 using Fragpipe^40^. The average of all precursor RAAS values from human tissues was taken as a point of comparison to RAAS values measured from IP-MS experiments.

### Ionization efficiency estimation

Ionization efficiencies of peptides were computed according to the model published by Liigand, *et al.* [33]. The authors established that for small peptides, such as those produced by trypsin digestion, the ionization efficiencies can be approximated as the sum of the ionization efficiencies of the amino acids comprising the peptide. They also provided empirical measures of the ionization efficiency of each amino acid. Therefore, the ionization efficiency of a SAAP or BP is estimated by the sum of the ionization efficiencies of the amino acids that make up the peptide.

### Peptide detectability prediction

Peptide detectability was computed using the DeepMSPeptide method described by Serrano, *et al.*^34^. The script DeepMSPeptide.py was downloaded from the DeepMSPeptide project github and was executed with a list of substituted and base peptide sequences as input. The resulting detectability scores were compared between substituted and base peptides.

### Evaluating the internal consistency of RAAS estimates

To test the reliability of high RAAS, we evaluated their consistency with the correlation patterns of BP or SAAP abundance to shared peptide abundance across patients, Extended Data Fig. 3i. This test capitalized on the internal consistency of different peptides from the encoded (P1) and alternatively translated (P2) proteoforms, thus allowing triangulation across multiple measurements, Fig. 2. Yet, this power is diminished by the uncertainty of proteoform composition and thus the shared peptide associated with each pair of BP and SAAP. This correlation analysis used the intensities of sample-specific reporter ions that are unlikely to share acquisition artifacts with the precursor intensities used by our consistency test described in Fig. 2d. The data indicate that as RAAS increases, suggesting reduced relative proportion of the P1 proteoforms, the correlations between BP and shared peptides, CORR(BP,shared peptides), significantly decrease, Extended Data Fig. 3i (top panel). Meanwhile, the correlations between SAAP and shared peptides, CORR(SAAP,shared peptides), increase slightly for peptides with RAAS*>*0, Extended Data Fig. 3i (middle panel) and the difference between CORR(SAAP, shared peptides) and CORR(BP, shared peptides) increases with increasing substitution ratios, Extended Data Fig. 3i (bottom panel). Many SAAP with high substitution ratios correlate to shared peptides more strongly than their counterpart BP, an effect most significant for peptides with RAAS*>*1 (Fig. 3i). This trend indicates that our RAAS estimates are consistent with triangulated measurements across shared peptides, despite the caveat of uncertain proteoform structures.

### Validation in multiple digests

To establish confidence in the data and evaluate potential impacts of the protease used for protein digestion, we used the tonsil sample from the label-free dataset^26^, which the authors subjected to digestion by several proteases. Each digest was processed through the SAAP identification, validation and quantification pipeline described above. Substituted peptides found in each digest were mapped back to their protein to match substitution sites across digests and compute the percentage of substitutions identified in multiple digests. To determine the percentage of base peptides identified in multiple digests, we chose a random amino acid from the tryptic base peptide to search whether it is detected as part of a peptide identified from the alternative proteases.

### SAAP degradation rates

Raw LC-MS proteomics data from SILAC-labeled human cell lines ref.^39^ was downloaded from the publication data repository (B cells: PXD008511, hepatocytes: PXD008512, monocytes: PXD008513, NK cells: PXD008515) and processed through the SAAP identification pipeline described above. Since there was no corresponding transcriptomic data, the DP search in MaxQuant was conducted with the human protein sequence fasta, downloaded from Uniprot. Lys8 and Arg10 were included as additional variable modifications to account for the SILAC-introduced heavy labels. After identifying candidate SAAP, these sequences were appended to the Uniprot database, along with the validated SAAP from the CPTAC and label-free^26^ datasets. To increase confidence in our results by using an independent search engine, the validation search was done using MS-Fragger^40^ with the SILAC3 workflow and MSBooster^41^ enabled. All identified heavy and light peptides, including validated SAAP, BP and other peptides, were quantified by summing the precursor ion intensity over all charge states and modifications (other than heavy isotopes) and taking the median across the replicates. RAAS was calculated as described above. Degradation rates (*α*) were computed using linear regression as the slope of the ln(1 + *h/l*) = *αt*, where *h/l* is the ratio between the heavy (h) and light (l) peptides abundance across time.

### Protein set enrichment analysis

Protein RAAS values were computed as the median RAAS of all SAAP mapping to the protein. Gene ontology (GO, geneontology.org) biological processes enriched with high RAAS were identified by comparing the distribution of RAAS for proteins in a given biological process to the distribution of RAAS for all substituted proteins using a Kolmogorov-Smirnov test to obtain p-values, which were then adjusted with a Benjamini-Hochberg FDR correction. GO processes with identical sets of substituted proteins were manually combined. Proteins representing significant biological processes are highlighted in Fig. 2g.

### Estimation of substituted protein copy number

Substituted protein copy numbers were estimated using the histone ruler approach introduced in ref.^32^. Briefly, the abundance of histones was computed as the median intensity of peptides corresponding to core histone proteins (H2A, H2B, H3, H4). This value was assumed to represent 30e6 protein copies, the approximate number of histone protein copies in a cell. SAAP protein copy numbers were estimated by multiplying the SAAP abundance, computed from precursor ion intensities as described above, by 30e6/histone protein abundance.

### RAAS dotplots

The global distribution of log_10_(RAAS) is approximately log-normal. Therefore, the distributions for various subsets of the observed substitutions (substitutions corresponding to a specific codon, substitution type or amino acid property, all substitutions in a given protein or a GOslim annotated group of proteins, and substitutions classified by disorder and conservation score bins), were compared to the distribution of all other values from the same distribution (globally for Extended Data Fig. 4b,d, Fig. 5c and Extended Data Fig. 5a, or within cancer types for Fig. 3b,c,e, Fig. 4c, Extended Data Fig. 4c and Extended Data Fig. 5c,f) by two-sided Student’s t-tests (R’s *t.test* function with the Welch approximation, called from our custom function `raasProfile`). The p-value, sidedness (test statistic *t <* 0 or *t >* 0) and log_10_ of the median RAAS value were recorded, and used to generate the RAAS dotplots (function `dotprofile`), where the dot size scales with − log_10_(*p*) up to a cut-off, and dot colors scale with the median RAAS value, as indicated by individual Figure legends. The number of RAAS values that were considered for each row and column group are indicated on the top and right axes. RAAS dotplots in the main manuscript often show only a subset of the analyzed categories filtered for having at least one significantly high or low RAAS distribution, as indicated in the Figure captions, and using the `sortOverlaps` function from the segmenTools R package (https://github.com/raim/segmenTools/ release `RAAS preprint`).

### Correlation of RAAS profiles

Median RAAS across all SAAP with a given substitution type was computed for each substitution type in each dataset, a subset of which is displayed in Fig. 3b. Pearson correlations weighted by the number of SAAP associated with each substitution type were computed using vectors of these values for every pair of datasets. The resulting correlation heatmap of RAAS profiles for substitution types across datasets is shown in Fig. 2d. Unweighted Pearson correlations across datasets are similarly computed for RAAS profiles of encoded and incorporated amino acid types, as displayed in Extended Data Fig. 5f,g.

### ANOVA analyses

To increase our confidence that variability in RAAS is biologically driven as opposed to driven by technical differences between datasets, we fit an ordinary least squares multiple regression model with precursor-level RAAS values for all SAAP in all datasets as the dependent variable and encoded amino acid type, incorporated amino acid type, substitution type, tissue type, and dataset type (TMT or label-free) as the independent variables. The contribution of each of the model variables to variance in RAAS was determined by an analysis of variance (ANOVA) test. The F-value, representing the fraction of explained variance, and p-value for each feature is displayed in Fig. 3f. A similar analysis was executed to determine the contribution of species (human/mouse) differences to RAAS variance relative to the contributions of substitution and tissue types, Fig. 5f.

### Substitution enrichment in tissue types

To determine enrichment of substitution types in specific tissues, we focused on tissues with both CPTAC and label-free healthy data, namely kidney (CCRCC), pancreas (PDAC), lung (LUAD, LSCC) and endometrium (UCEC). For each substitution type, we computed the log_2_(fold change) and p-value (t-test) between RAAS of all SAAP with that substitution type identified in a given tissue and the RAAS of all SAAP with that substitution type measured in all other tissues. p-values were adjusted with a Benjamini-Hochberg FDR correction to get q-values. Substitution types were considered significantly enriched with high or low RAAS values in a given tissue if the log_2_(fold change) relative to all other tissues was *>* 1 or *<* −1, respectively, with a q-value of ≤ 0.01. Substitution types with significantly high RAAS in a given tissue are highlighted in Extended Data Fig. 5h,i.

### Comparison of substitution type RAAS in tumor and normal tissue

Log_2_ fold changes of RAAS between tumor and matched normal tissue for a given substitution type was computed by taking the mean of the log_2_ ratios of the RAAS for each SAAP with the substitution type in a patient tumor sample relative to the RAAS for the same SAAP in the patient-matched normal adjacent tissue sample. p-values were computed with a paired t-test. p-values were FDR-adjusted into q-values using the Benjamini-Hochberg method.

### Pfam domain analysis

The canonical isoforms of proteins with substitutions were mapped to Pfam structural protein domains using InterProScan v5.67-99.0, which was downloaded from EMBL-EBI (InterProScan) and run on the Discovery high performance computing cluster (MGHPCC, Holyoke, MA). The significance of finding amino acid substitutions in a given Pfam domain was computed by counting the number of substitutions identified in a domain and comparing to a bootstrapped distribution of substitution counts in the domain obtained by shuffling the position of the substitutions across the entire protein sequence. The p-value was computed as the probability of finding more substitutions in the domain than expected by chance (random shuffling of substitutions). Domains with significantly high or low RAAS were identified by comparing the distribution of RAAS for substitutions in a domain to the distribution of all RAAS values using a Kolmogorov-Smirnov test and Benjamini-Hochberg FDR correction. Pfam domains significantly enriched with substitutions of significantly high or low RAAS are displayed in Extended Data Fig. 6d.

### Substitution sequence context analysis

To analyze the sequence context of amino acid substitution sites, we generated a custom protein database of all proteins defined in the Ensembl human genome release GRCh38.110, supplemented with all proteins with patient-specific mutations from the sample-specific protein database generation pipeline described above. All unique base peptides were searched against this database using blastp (NCBI’s BLAST+, version 2.15.0+), and only blast hits with 100% sequence identity and no mismatches were considered. For these hits, we obtained the amino acid context (sequences surrounding the substitution sites) from the original Ensembl protein and the codon from the matched Ensembl transcript. If the substitution itself covered a patient-specific mutation the codon was not considered. GOslim annotations for all genes were downloaded via Ensembl BioMart queries on 2023-12-04. Relative codon frequencies were calculated from the full Ensembl transcripts of the set of all Ensembl proteins with identified AAS sites. The total count for each codon was divided by the total count of all codons for the same amino acid.

### Sequence Difference Logos

Difference logos were generated with the R package DiffLogo^78^ (version 2.26.0) using a customized DiffLogo function that allows for custom y-axis limits of the plots; and p-values were calculated with the packages’ `enrichDiffLogoObjectWithPvalues` function, but are only shown for residues that were not part of the sequence selection (e.g. the substitution site when selecting sequences by encoded and incorporated amino acid). In total 7,086 unique sequences surrounding the substitution sites were considered, and for all difference logos, a selected subset of these sequences was compared to all other sequences.

### Sorted enrichment profiles

Amino acid enrichment around unique substitution sites (Extended Data Fig. 8a,b) and the location of substitution sites along base peptides (Extended Data Fig. 8c) were evaluated using functions of the R package segmenTools, as described in detail by Behle et al.^79^. Shortly: enrichments were calculated by cumulative hypergeometric distribution tests (function `clusterCluster`). Significantly enriched (*p* ≤ 10*^−^*^5^) amino acids or sites were sorted along the x-axis from top to bottom (function `sortOverlaps`). The field color scales with −*log*_2_(*p*), cut at *p* ≤ 10*^−^*^10^ and the white text indicates *p* ≤ 10*^−^*^5^ (function `plotOverlaps`).

### Protein structural features

IUPred3^80^ was used to calculate a “disorder” score for each protein in our database, with options `-a` to add the `ANCHOR2` prediction of disordered binding sites, and `-s medium` to use the “medium” smoothing type. Pfam domains for each protein were predicted with HMMER3^81^ (version 3.4 (Aug. 2023)), using a reporting cutoff `-E 0.01` for a comprehensive output. Protein secondary structures were predicted by S4pred^82^ (version 1.2.5, downloaded from https://github.com/psipred/s4pred on 2024-01-26). Alternative scores for disordered (flDPnn^83^ and disordered protein interaction sites (disordRDPbind^84^), as well as a MMseqs2^85^-based sequence conservation score were obtained *via* the DescribePROT database^86^. Download and calculation of protein structural features is documented at https://gitlab.com/raim/genomeBrowser/-/tree/master/data/mammary (release `RAAS preprint`). ChimeraX^87^ was used to display selected PDB structures and color substituted sites by their median RAAS value.

### Predicting RAAS from sequence features

To understand how much of the site to site variance on RAAS could be explained by known protein, local sequence, and amino acid site specific structural properties, we created an classifier trained on a variety of features to predict RAAS. XGboost^88^, R implementation version 1.7.7.1, was chosen for the classifier because it allowed for easily incorporating missing data features that were not completely known across the entire proteome. The data was randomly split into a train fraction, 80 %, and a test fraction, 20 % to evaluate model. The model was retrained 1000 times on random subsets of test and train data. The scatter plot was a representative sample reflecting the median correlation between train and test. The model was then retrained 100 times in the same manner leaving out one factor each time to estimate the added predictive power for each feature, defined as the decrease in correlation from the all feature model.

### Allele frequency of missense variants in alternatively translated codons

Allele frequencies in gnomAD v4.1.0 (730,947 exomes)^49^ were calculated for two sets of missense substitutions: (1) all possible single-nucleotide missense variants in codons affected by alternative translation (Fig. 5e), and (2) single-nucleotide mutations in the codon that result in the alternate amino acid when translated according to the genetic code (Extended Data Fig. 10d). Codons were grouped into quartiles based on their RAAS values. The site frequency spectrum (SFS) for each group was binned into eight allele frequency categories: 0 (monomorphic), 0 to 10*^−^*^6^, 10*^−^*^6^ to 10*^−^*^5^, 10*^−^*^5^ to 10*^−^*^4^, 10*^−^*^4^ to 10*^−^*^3^, 10*^−^*^3^ to 10*^−^*^2^, 10*^−^*^2^ to 10*^−^*^1^, and 10*^−^*^1^ to 0.5.

### Missense variant constraint in alternatively translated codons

Constraint was calculated for three sets of missense substitutions in the gnomAD v4.1.0 dataset (730,947 exomes)^49^: (1) all possible single-nucleotide missense variants in codons affected by alternative translation (Fig. 5f), (2) single-nucleotide mutations that lead to the observed SAAP sequences when translated according to the genetic code (Extended Data Fig. 10e), and (3) single-nucleotide mutations that do not lead to the alternate amino acid in SAAP when translated according to the genetic code. For each of these sets, across all RAAS quartiles, constraint was measured by dividing the observed number of mutations by the expected number based on the Roulette mutational model^89^. This model predicts the expected number of mutations at each site by considering nucleotide context and known mutational processes in the human genome. Two control groups were used for comparison within each RAAS quartile. The first control group contained the most deleterious missense variants in genes from this RAAS quartile, as predicted by AlphaMissense^90^ (top decile). These variants had an observed/expected ratio close to 0.5 indicating strong constraint (twice as constrained as synonymous mutations). The second control group contained the least deleterious missense variants in genes from this RAAS quartile (lowest decile), with an observed/expected ratio near 1 indicating minimal constraint (no more constrained than synonymous mutations). AlphaMissense is a machine learning-based tool that predicts the deleteriousness of missense variants by integrating functional annotations and evolutionary conservation data. The use of these control groups allows for benchmarking the constraint observed in the sets of interest against variants with known levels of deleteriousness.

### Overlap with RNA modification sites

To explore whether uracil (U) modifications may contribute to alternate RNA decoding, we analyzed the nucleotide overlaps of codons corresponding to amino acid substitutions with sites of modified U residues in transcripts, as previously defined by nanopore sequencing^38^. We mapped all modified U sites from 6 cell lines to the coding sequences of the set of Ensembl MANE transcripts (Matched Annotation from NCBI and EBI), yielding 39,723 unique sites in 6,771 distinct transcripts. We also reduced the set of AAS from 7,069 to 6,967 sites that map to MANE transcripts. We then counted all U in the union of all 7,471 transcripts with AAS and/or *ψ* sites as the total set, and all 4,250 U in codons at AAS sites as the test set, and found 180 overlapping sites. To test whether this is a significant enrichment we used a cumulative hypergeometric distribution test (*p*[*X >* 179], in R: `phyper(q=180-1, m=39723, n=2879635-39723, k=4250, lower.tail=FALSE`). To further evaluate the calculated p-value, we plotted the p-value distribution over different counts in panel and indicate the real count and its associated p-value in red, Supplemental Fig. 8a.

## Availability

Supporting information, data, and documentation is available at decode.slavovlab.net. The code is freely available at github.com/SlavovLab/decode.

## Acknowledgments

We thank Dr. Mahlon Collins for help with analysis, Prof. Nuno Bandeira, Prof. Arjun Raj, Prof. Zoya Ignatova and Prof. Barry Karger for detailed feedback, Orhun Kok and Shanshan Zheng for help with the revision, and members of the Slavov laboratory for discussions and suggestions. The work was funded by an Allen Distinguished Investigator award through The Paul G. Allen Frontiers Group to N.S., an NIGMS award R01GM144967 to N.S., NCI awards UG3CA268117 and UH3CA268117 to N.S., an NIGMS award R35GM148218 to N.S., and an NIA award R01AG092460 to N.S., and a Bits to Bytes award from MLSC to N.S

## Competing interests

Nikolai Slavov is a founding director and CEO of Parallel Squared Technology Institute, which is a non-profit research institute.

## Author contributions

**Study design, supervision, and raising funding**: N.S.

**Data analysis**: S.T., R.M., A.L., S.W., E.K., and N.S.

**gnomAD analysis:** J.G., S.T., and K.K.

**Initial draft**: S.T., and N.S.

**Writing**: All authors approved the final manuscript.

## Abbreviations

SAAP: substituted amino acid peptide (product of alternate translation)
BP: base peptide translated according to the genetic code
RAAS: ratio of amino acid substituted peptide to the corresponding non-substituted peptide
CCRCC: clear cell renal cell carcinoma; BRCA – breast invasive carcinoma
UCEC: uterine corpus endometrial carcinoma
PDAC: pancreatic ductal adenocarcinoma
LUAD: lung adenocarcinoma
LSCC: lung squamous cell carcinoma

## Notes

https://decode.slavovlab.net/

https://github.com/SlavovLab/decode

### Summary of Updates

The previous version had lower quality figures due to file size limitations. This version has higher quality figures and the file size was reduced by placing the extended data figures and supplemental figures in the supporting information.

