## Extended Data Figures and Supplemental Figures for "Alternate RNA decoding results in stable and abundant proteins in mammals"

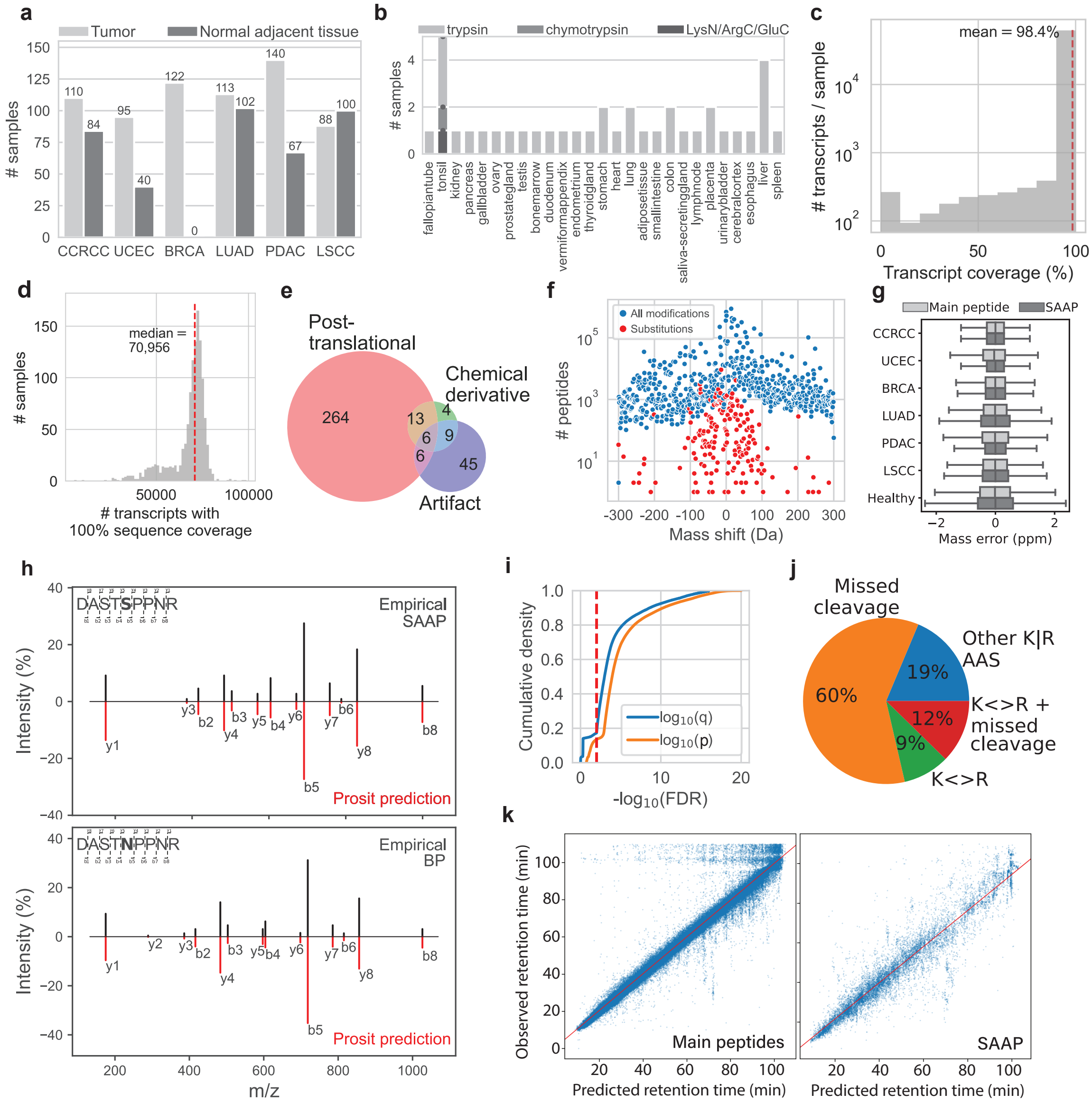


**Extended Data Fig. 1 | Systematic identification and validation of amino acid substitutions** (**a**) Number of tumor and normal samples analyzed from each CPTAC dataset. (**b**) Number of samples analyzed for each healthy tissue from the label-free dataset. (**c**) Distribution of the percentage of each transcript with a read that is included in the patient-specific databases. (**d**) Distribution of the number of transcripts with 100% sequence coverage included in each patient-specific protein database. (**e**) Non-substitution modifications identified in the dependent peptide search are majorly comprised of post-translational modifications, and include artifacts and chemical derivatives from MS analysis. (**f**)– (**k**) (Continued on the next page)

(Continued) (**f**) The number of modified peptides identified as having an amino acid substitution or other type of post-translational or chemical modification. (**g**) Mass error distributions for SAAP and all peptides identified in the database search show no significant differences. (**h**) Butterfly plots showing a systematic mass shift in MS2 spectra between SAAP and BP for a representative SAAP with median RAAS=1.2 in pericentriolar material 1 protein isoform 1 (PCM1). The fragmentation spectra were predicted by the Prosit TMT model[75]. (**i**) Cumulative density distributions of p-values (MaxQuant) and FDR-controlled q-values computed using only SAAP. Red dashed line indicates confidence threshold for SAAP inclusion in further analysis. (**j**) Over 80% of substitutions identified from lysine (K) or arginine (R) are at sites of missed cleavage or are substitutions between K and R. (**k**) Observed and predicted (DeepRT+,[28]) retention times show strong agreement for all main peptides identified in standard database search and for SAAP.

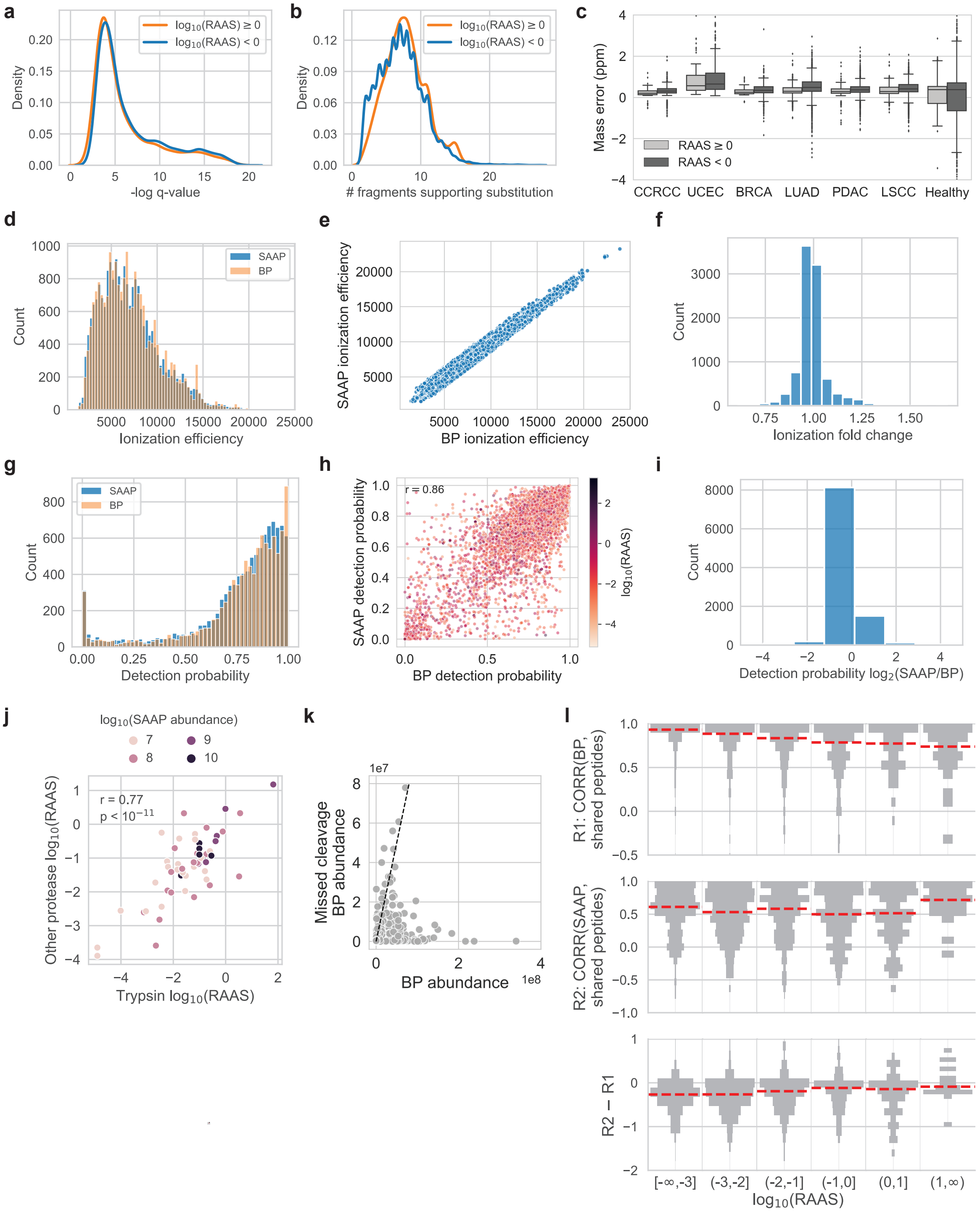


**Extended Data Fig. 2 | Establishing confidence in AAS abundance** (**a**) SAAP with high RAAS≥0 are identified with the same FDR-controlled confidence as SAAP with low RAAS<0. (**b**) SAAP with high RAAS≥0 are identified with as many fragment ions providing evidence at the site of alternate translation as SAAP with low RAAS<0. (**c**) SAAP with high RAAS≥0 have similar mass error distributions as SAAP with low RAAS<0. (**d**)– (**l**) (Continued on the next page)

(Continued) **(d-f)** SAAP abundance estimates are unlikely to be affected by differences in ionization efficiency between base peptides and alternatively translated peptides. The ionization efficiency distributions for SAAP and BP are indistinguishable (**d**), correlate strongly (**e**), and there is negligible fold change between them (**f**). **(g-i)** SAAP abundance estimates are unlikely to be affected by differences in peptide detectability between base peptides and alternatively translated peptides. The peptide detectability distributions for SAAP and BP are indistinguishable (**g**), correlate strongly (**h**), and there is negligible fold change between them (**i**). (**j**) SAAP abundances (RAAS) computed for the same site of alternate translation from peptides in different enzymatic digests of tonsil are consistent for peptides across the range of peptide abundances. (**k**) Base peptides with missed cleavages are generally an order of magnitude more lowly abundant than their fully cleaved counterparts. (**l**) Correlation of BP (top panel) or SAAP (middle panel) abundance with shared peptide abundance across samples. Shared peptides are peptides found in both the encoded and alternatively translated proteoforms. BP correlation to shared peptides decreases with increasing RAAS, while SAAP correlation to shared peptides tends to increase, especially at RAAS$>$1, as indicated by the difference in these correlations (bottom panel), and in support of the hypothesis presented in Fig. 2a. Abundances are computed with MS2-level intensities.

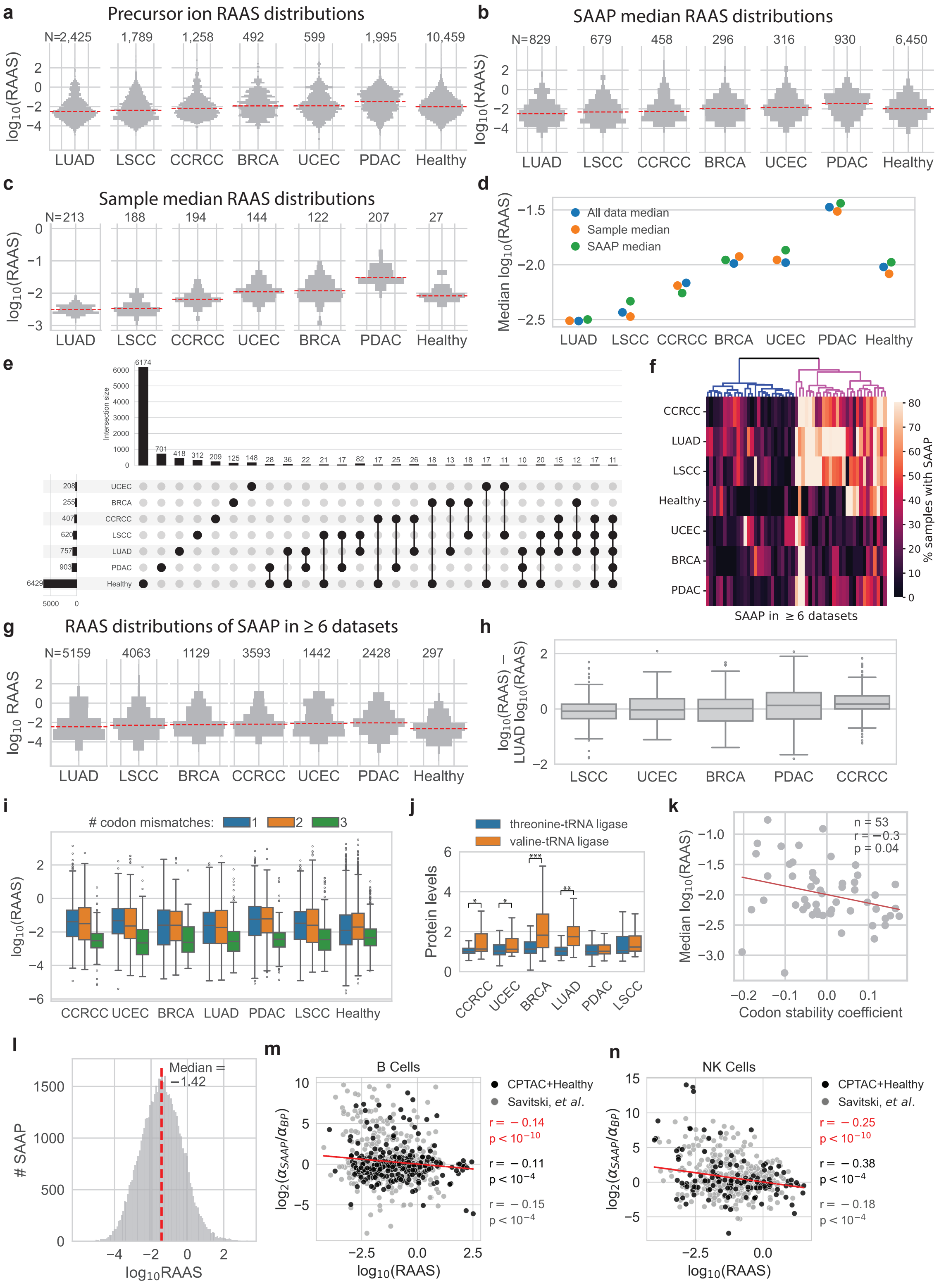


**Extended Data Fig. 3 | Quantification of substituted amino acid peptides (SAAP) (a)– (k)** (Continued on the next page)

(Continued) (**a**) Distributions of substitution ratios for all SAAP identified in each dataset computed for each experiment (TMT set, CPTAC data) or sample (label-free data), using MS1 precursor ion intensities. N indicates the number of RAAS computed at the MS1 level in each dataset. (**b**) Median RAAS were computed for each unique SAAP-BP pair using reporter ion intensities (MS2, CPTAC data) or precursor ion intensities (MS1, label-free data). Distributions of median RAAS across all SAAP in a dataset are shown. N indicates number of unique SAAP-BP pairs identified in each dataset. (**c**) Median RAAS across all SAAP identified in each sample were computed using reporter ion intensities (MS2, CPTAC data) or precursor ion intensities (MS1, label-free data). Distributions shown are of these medians across all samples in a dataset. N indicates the number of samples in each dataset. (**d**) Substitution ratio distributions shown in (**a**), (**b**), (**c**) have consistent medians, highlighting variability in RAAS across datasets. (**e**) Upset plot showing overlap in unique SAAP identified across all datasets. Dataset combinations require at least 10 shared SAAP to be included in visualization. (**f**) Heatmap displaying the percentage of samples in each dataset in which SAAP identified in 6+ datasets are found. Hierarchical clustering shows a cluster of shared SAAP that are commonly identified across majority of samples in addition to 6+ datasets. (**g**) To confirm variability in RAAS across datasets, we looked at the subset of SAAP that were identified in at least 1 sample in at least 6 datasets. LUAD and LSCC substitutions consistently have the lowest RAAS, while PDAC substitutions have the highest RAAS. N indicated the number of RAAS computed for shared SAAP in each dataset. (**h**) Boxplots highlighting the difference between RAAS in CPTAC datasets relative to RAAS computed in LUAD. Only SAAP shared between LUAD and the compared dataset are used. Each data point is a $\log_{10}$(RAAS) difference computed for a unique SAAP-BP pair. (**i**) RAAS as a function of the minimum number of codon-anticodon mismatches needed for incorporating the detected amino acid across all datasets. (**j**) An example of a substitution that can be partially explained by synthesis errors arising from significantly (t-test) higher abundance of the amino acyl-tRNA ligase supplying the alternatively translated amino acid relative to the abundance of the amino acyl-tRNA ligase supplying the encoded amino acid. *: q-value$< 10^{-3}$, **: q-value$< 10^{-5}$, ***: q-value$< 10^{-20}$. (**k**) RAAS negatively correlates to the codon stability coefficient, an empirical measure of codon usage. n denotes number of codons, r is Pearson correlation, p is correlation p-value, and the red line is the ordinary least squares fit. (**l**) RAAS distributions for SAAP identified and validated in human hepatocytes. (**m**) The stability of SAAP relative to BP in primary human B cells is inversely proportional to their RAAS (Pearson correlation). (**n**) Same as (**m**) but in NK cells.

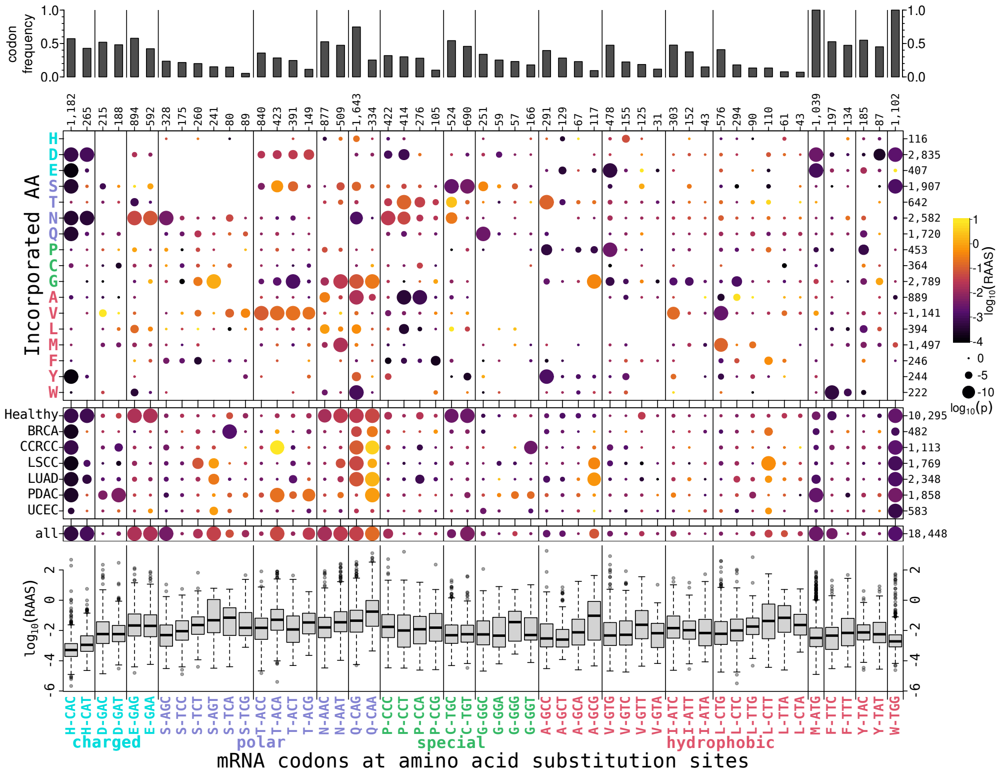


**Extended Data Fig. 4 | Associations between codons, incorporated amino acids and RAAS** (**a**) Relative codon frequencies in the full transcripts of the set of all proteins with identified substitution sites. The total count for each codon was divided by the total count of all codons for the same amino acid. Codon groups (per amino acid, separated by vertical lines) were sorted by amino acid property groups. Within each codon group, codons were sorted by their relative frequencies. All other panels are aligned with this sorting, see (**e**) for x-axis labels. These relative frequencies (bar heights) were also used in Fig. 2j (x axis). (**b**) RAAS dotplot for codons (columns) and incorporated amino acids (rows), sorted and color-coded by amino acid property groups. (**c**) RAAS dotplot for codons by datasets, i.e., cancer types and healthy tissues. (**d**) RAAS dotplot for all codons without further subsetting. Note, that the median RAAS values (colors) correspond to the y-axis values in Fig. 2j. (**e**) RAAS distributions for each codon.

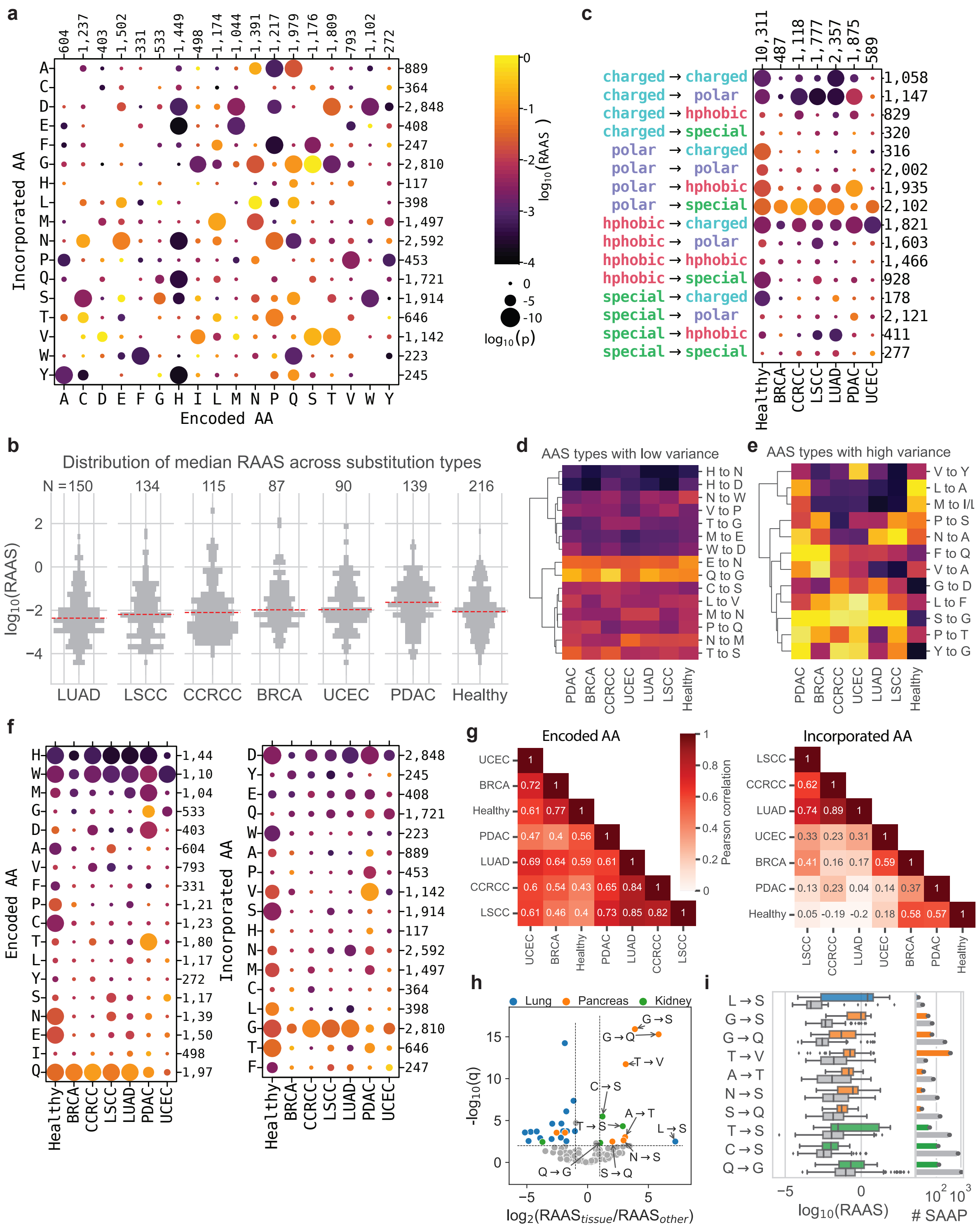


**Extended Data Fig. 5 | Substitution ratios depend on substitution and tissue types** (**a**) RAAS dotplot for all encoded and incorporated amino acids. (**b**)– (**h**) (Continued on the next page)

(Continued) (**b**) Violin plots of RAAS medians for each substitution type in every dataset. N indicates number of substitution identified in each dataset. (**c**) RAAS dotplot as in (**a**) but by chemical properties of the encoded and incorporated amino acids. (**d**) Heatmap of median RAAS by substitution type for substitution types with variance $<10\%$ across datasets. (**e**) Heatmap of median RAAS by substitution type for substitution types with variance $>50\%$ across datasets. (**f**) RAAS dotplots for encoded (left panel) and incorporated (right panel) amino acids. (**g**) Median RAAS values for SAAP grouped based on the encoded amino acid (left panel) or incorporated amino acid (right panel) correlate strongly and significantly across all datasets (Pearson correlation). (**h**) Substitution types with significantly higher RAAS in a given tissue type (cancer and healthy samples) relative to all other tissues analyzed (t-test, Benjamini-Hochberg FDR-corrected). (**i**) RAAS distributions (boxplots) and number of SAAP identified (barplot) for substitution types that are significantly higher in a given tissue (colored) relative to all other tissues (gray) analyzed. Colors indicate the same tissue types as in (**h**).

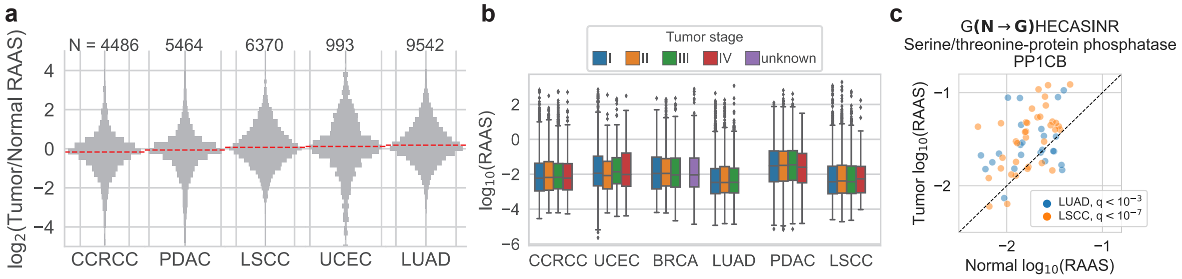


**Extended Data Fig. 6 | Associations of substitutions with cancer** (**a**) RAAS fold change between tumor and patient-matched normal adjacent tissue samples with median of distribution shown in red. N indicates the number of patient-specific RAAS values compared. (**b**) Patient-level RAAS distributions stratified by clinical tumor stage. No significant associations were measured between RAAS and tumor stage. (**c**) RAAS for the $N \rightarrow G$ substitution in serine/threonine-protein phosphatase PP1-beta catalytic subunit (PP1CB) is significantly higher in tumor samples than in patient-matched normal adjacent tissue, for the majority of patients in LUAD and LSCC (t-test).

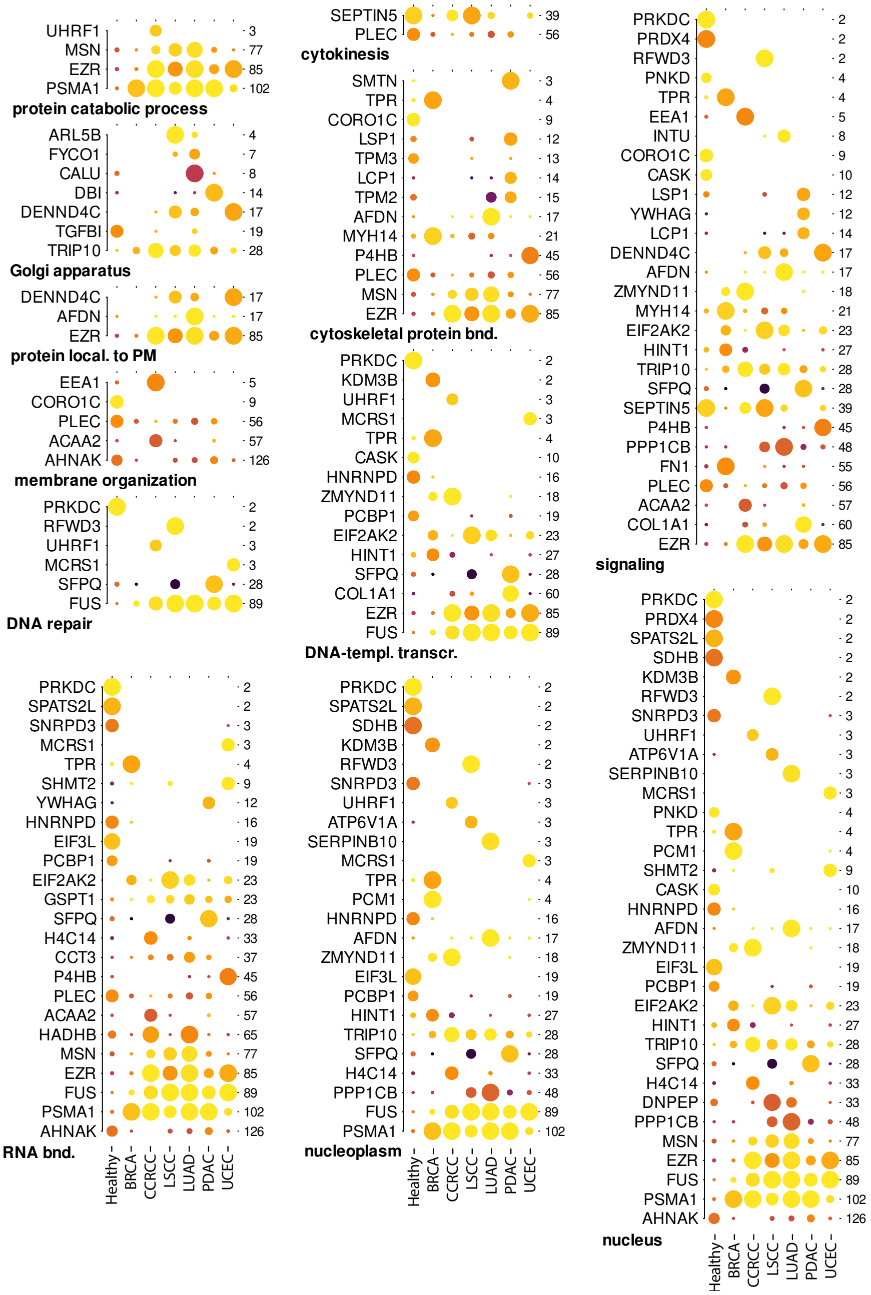


**Extended Data Fig. 7 | Proteins with high RAAS organized by functional groups** RAAS dotplots for all proteins with significantly high median RAAS ($p \leq 10^{-5}$) from each functional category shown in Fig. 4c.

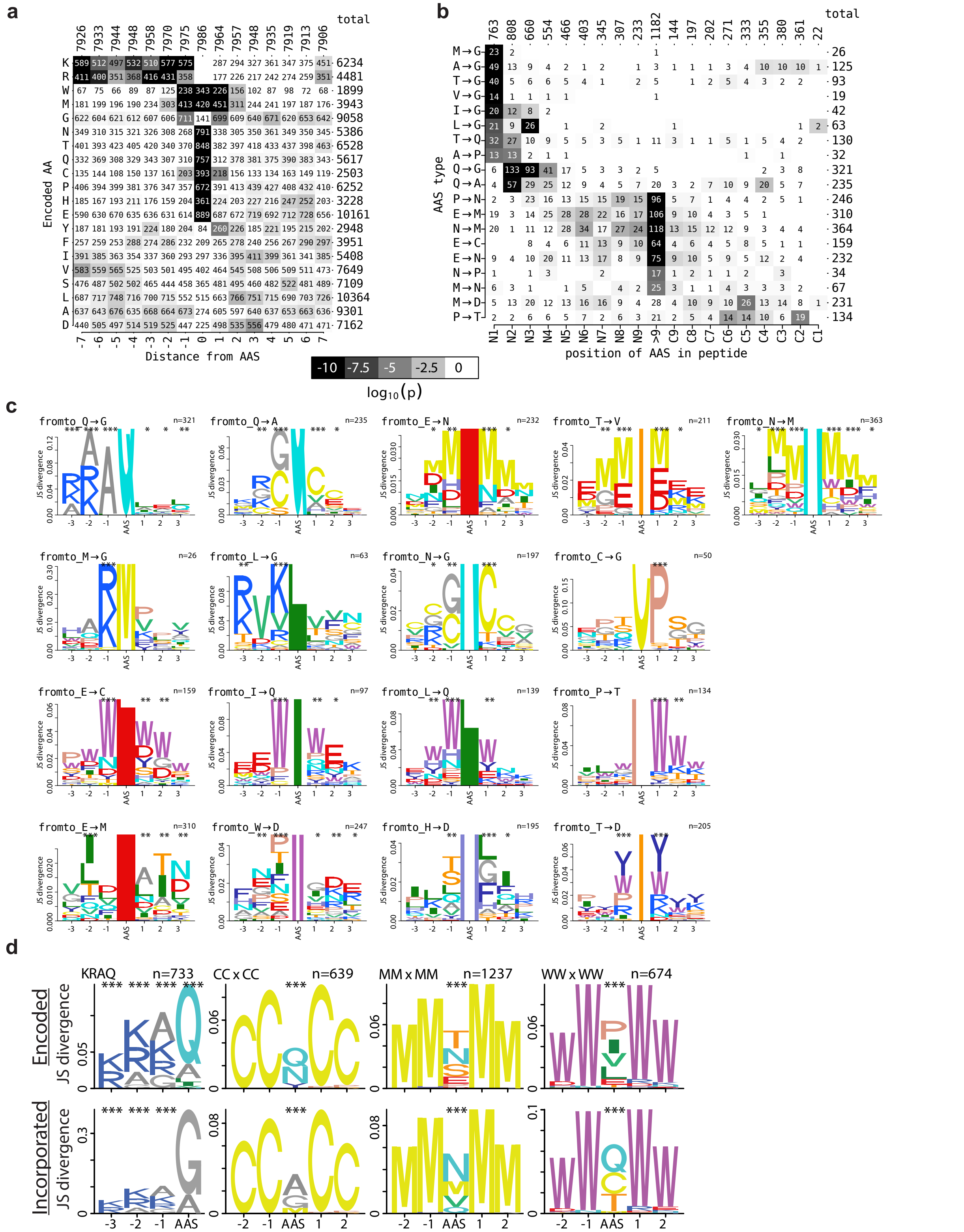


**Extended Data Fig. 8 | Amino acid sequence context around substitution sites**(Continued on the next page)

(**a**) Counts (text) and enrichments (gray scale: p-values of cumulative hypergeometric distribution tests) of amino acids surrounding the amino acid substitution sites. Lysine (K) and arginine (R) are enriched directly upstream of the substitution sites. Tryptophan (W), methionine (M), glycine (G) and cysteine (C) are enriched directly adjacent to substitution sites. (**b**) As (**a**) but for substitution types (encoded:incorporated) vs. their position in identified peptides. Substitutions by glycine ($\rightarrow$G) or alanine ($\rightarrow$A) are enriched within the 3 N-terminal amino acids of base peptides (N1 to N3), i.e., directly after the trypsin cleavage sites (K or R). Various substitutions involving arginine (N), methionine (M) or glutamate (E) as either the encoded or the incorporated amino acid are enriched distant from the N- and C-termini (>9). Only substitution types with at least one significant enrichment ($p \leq 10^{-10}$) are shown in (b). (**c**) Sequence difference logos were calculated for all unique sequences surrounding substitution sites, subset for all observed substitution types (encoded$\rightarrow$incorporated amino acids), and plots were only generated if any of the positions -3 to +3 around a substitution site showed a significant enrichment with $p \leq 10^{-10}$ (*: $p \leq 10^{-3}$, **: $p \leq 10^{-5}$, ***: $p \leq 10^{-10}$), and all resulting logos are shown. The logos were grouped by common patterns (rows from top to bottom): (i) Substitutions by glycine or alanine (Q$\rightarrow$A, Q$\rightarrow$G, M$\rightarrow$G, L$\rightarrow$G) are enriched directly upstream with lysine (K) or arginine (R), i.e. they are preferentially observed at the N-terminus of base peptides, next to the trypsin cleavage sites (K or R). (ii) Substitutions of glutamine (Q$\rightarrow$A) or arginine (N$\rightarrow$G) are flanked by cysteine (C) enrichments. (iii) substitutions E$\rightarrow$N, T$\rightarrow$V and N$\rightarrow$M are flanked by methionine (M) enrichments. (iv) Substitutions E$\rightarrow$C, I$\rightarrow$Q, L$\rightarrow$Q and P$\rightarrow$T are flanked by tryptophan (W) enrichments. (**d**) Sequence difference logos of selected subsets of substitution sites. AAS denotes the site of the substitution and numbers refer to adjacent positions in the protein sequence. The y-axis shows the Jensen-Shannon divergence of the selected set of sequences (number n of sequences is indicated) compared to all other sequences in our data; *** indicates enrichment significance $p < 10^{-10}$.

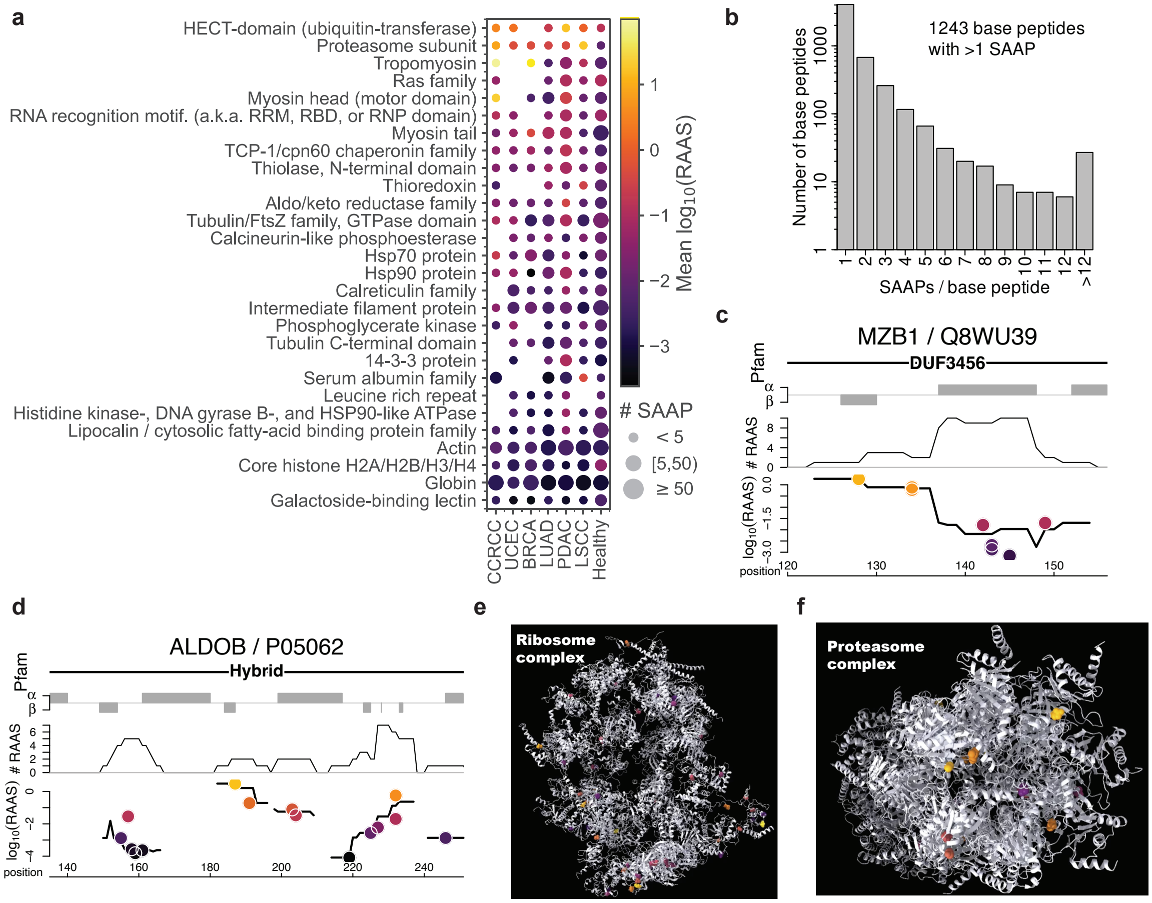


**Extended Data Fig. 9 | Substitution proximity in domains, 1D and 3D protein structures** (**a**) Pfam domains significantly enriched in substituted peptides (FDR-adjusted p-values $< 0.05$, see Methods for details). (**b**) The number of base peptides decreases exponentially with the number of distinct SAAP detected per base peptide. This reflects the abundance bias of AAS detectability, such that many distinct SAAPs can be detected for highly abundant peptides. (**c**) High density region of substitutions in marginal zone B- and B1-cell specific protein (MZB1). A high RAAS substitutions cluster in $\beta$-sheet and unstructured regions is immediately followed by a lower RAAS cluster in an $\alpha$-helix region. (**d**) Three 1D clusters of substitutions are found in the glycolytic region of aldolase B (ALDOB). (**e**) Many substitutions are identified in the ribosome complex, some of which cluster across complex subunits in the 3D protein structure. (**f**) High and low RAAS substitutions cluster in the 3D protein structure of the proteasome complex.

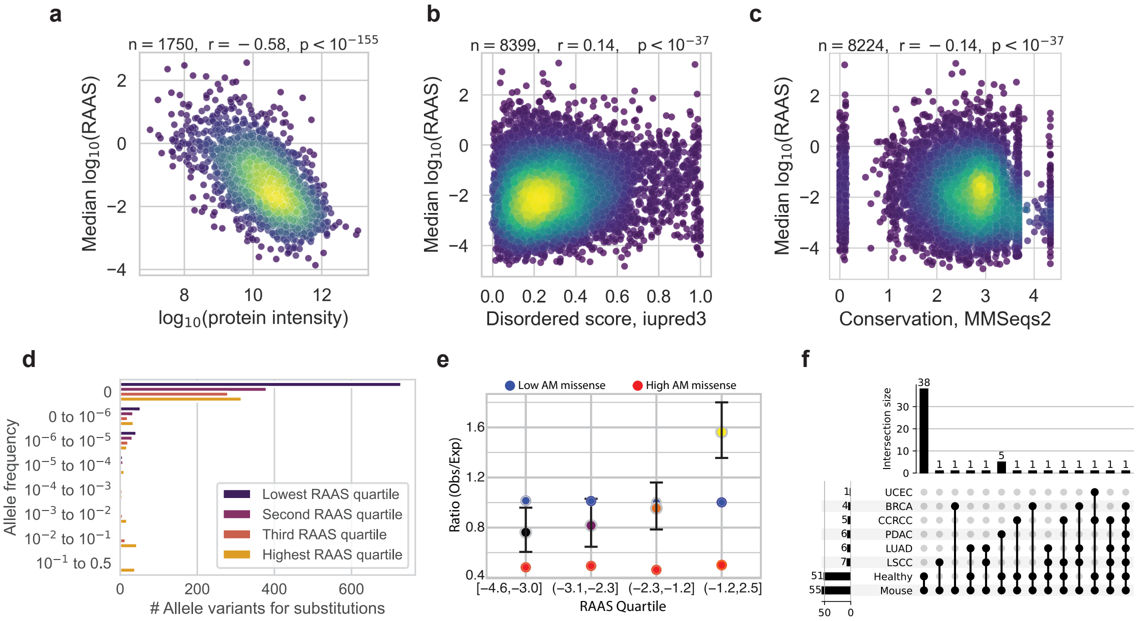


**Extended Data Fig. 10 | Associations of substitutions with protein properties** (**a**) Protein RAAS is negatively correlated to protein abundance. The intensity was calculated as the median over all "leading razor protein" intensities of all data sets (CPTAC and tissues) where a given base peptide was identified. (**b**) RAAS is positively correlated to the disordered score (IUPred3 prediction). RAAS was computed as the median RAAS for each unique BP/SAAP pair across all datasets (CPTAC and tissues) and the disordered score is the score of the protein at the AAS site. (**c**) as (**b**) but for the mean conservation score (MMseqs2 score via the DescribePROT database). (**d**) Allele frequency in the gnomAD database for missense variants coding for identified substitution in alternatively translated codons. Sites with allele variation frequency $\geq 10^{-3}$ correspond to 131 SAAPs. (**e**) Observed / expected ratios for missense variants coding for identified substitution in alternatively translated codons, determined from analysis of gnomAD database (see Methods). Missense variants are less constrained with increasing RAAS ($p < 10^{-6}$). Data point colors correspond to RAAS quartiles as in (**d**) or low or high AlphaMissense (AM) controls. (**f**) Upset plot showing overlap in unique SAAP identified between human and mouse tissues.

### Supplemental Information

#### Description of Supplemental Data tables

**Supplemental_Data_1.PTMs.csv** A table of peptides identified through the dependent peptide search as having known post-translational or chemical modifications. This table includes peptides from all 6 CPTAC datasets and the label-free healthy human tissue dataset. Types and locations of modifications are specified for each peptide.

**Supplemental_Data_2.SAAP_proteins.xlsx** A table containing 1 row per unique SAAP-BP pair per dataset (CPTAC and label-free). Each SAAP-BP pair is listed along with RAAS summary stats across the dataset and protein mapping information, including Pfam domains. The first tab of the file provides a description of the columns in the data table.

**Supplemental_Data_3.SAAP_precursor_quant.xlsx** A table containing 1 row per unique SAAP-BP pair per TMT set (CPTAC) or healthy tissue type (label-free). Each SAAP-BP pair is listed along with their precursor ion level abundances and RAAS values computed from precursor ion intensities. The first tab of the file provides a description of the columns in the data table.

**Supplemental_Data_4.SAAP_reporter_quant.xlsx** A table containing 1 row per unique SAAP-BP pair per patient sample (CPTAC only). Each SAAP-BP pair is listed along with their reporter ion level abundances and RAAS values computed from reporter ion intensities. The first tab of the file provides a description of the columns in the data table.

**Supplemental_Data_5.SAAP_SILAC_quant.xlsx** Table of SAAP-BP pairs identified in SILAC-labeled liver cells[39] containing one row per unique SAAP-BP pair per sample. Precursor ion level abundances of the light and heavy peptides in each sample are listed, along with their sums, ratios and RAAS values. The first tab of the file provides a description of the columns in the data table.

**Supplemental_Data_6.SAAP_neurodegenerative.xlsx** A subset of the substitutions from

Supplemental_Data_2.SAAP_proteins.xlsx corresponding to SAAP, BP and RAAS data pertaining to proteins that have been implicated in neurodegeneration and dementia (Uniprot).

**Supplemental_Data_7.SAAP_coordinates.tsv** Mapping of the amino acid substitution sites of unique pairs of BP/SAAP to proteins, their transcripts and genome coordinates, as defined in the Ensembl genome release `GRCh38.110`.

**Supplemental_Data_8.High_confidence_SAAP_precursor_quant.xlsx** Supplemental_Data_3 filtered for SAAP with and positional probability $> 0.9$.

**Supplemental_Data_9.gnomAD.xlsx** Data tables with gene locus, allele frequency and constraint data for all observed substitutions.

**Supplemental_Data_10.Aligned_reads.txt** Aligned reads to the transcripts coding for the base peptides (and corresponding SAAP) shown in Supplemental Fig. 1, 2 and 3.

**Supplemental_Data_11.IP-MS_SAAP.xlsx** SAAP identified in IP-MS pulldown experiments.

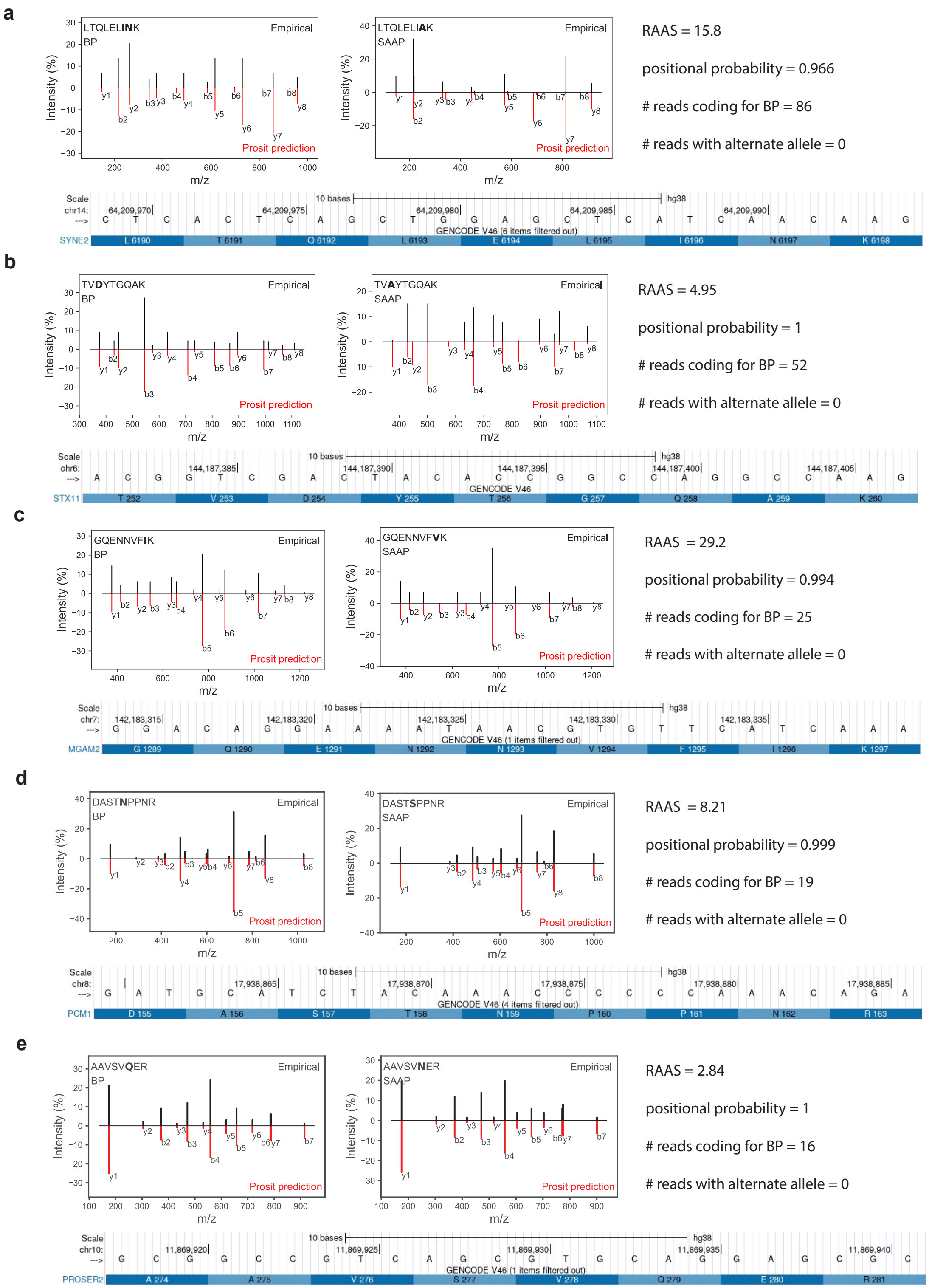


**Supplemental Fig. 1 | RNA-seq and fragment ion evidence for SAAP with RAAS>1 (part I).** Mirror plots display the empirical and predicted fragment ion spectra for 1 label-free (**a**) and 4 TMT-labeled SAAPs (**b**-**e**) and their corresponding BP, with substitution sites bolded in the peptide sequences. Genome browser tracks display the DNA sequence corresponding to the detected RNA molecules and the corresponding amino acid sequence predicted by the genetic code. The aligned reads are available in Supplemental_Data_10.Aligned_reads.txt.

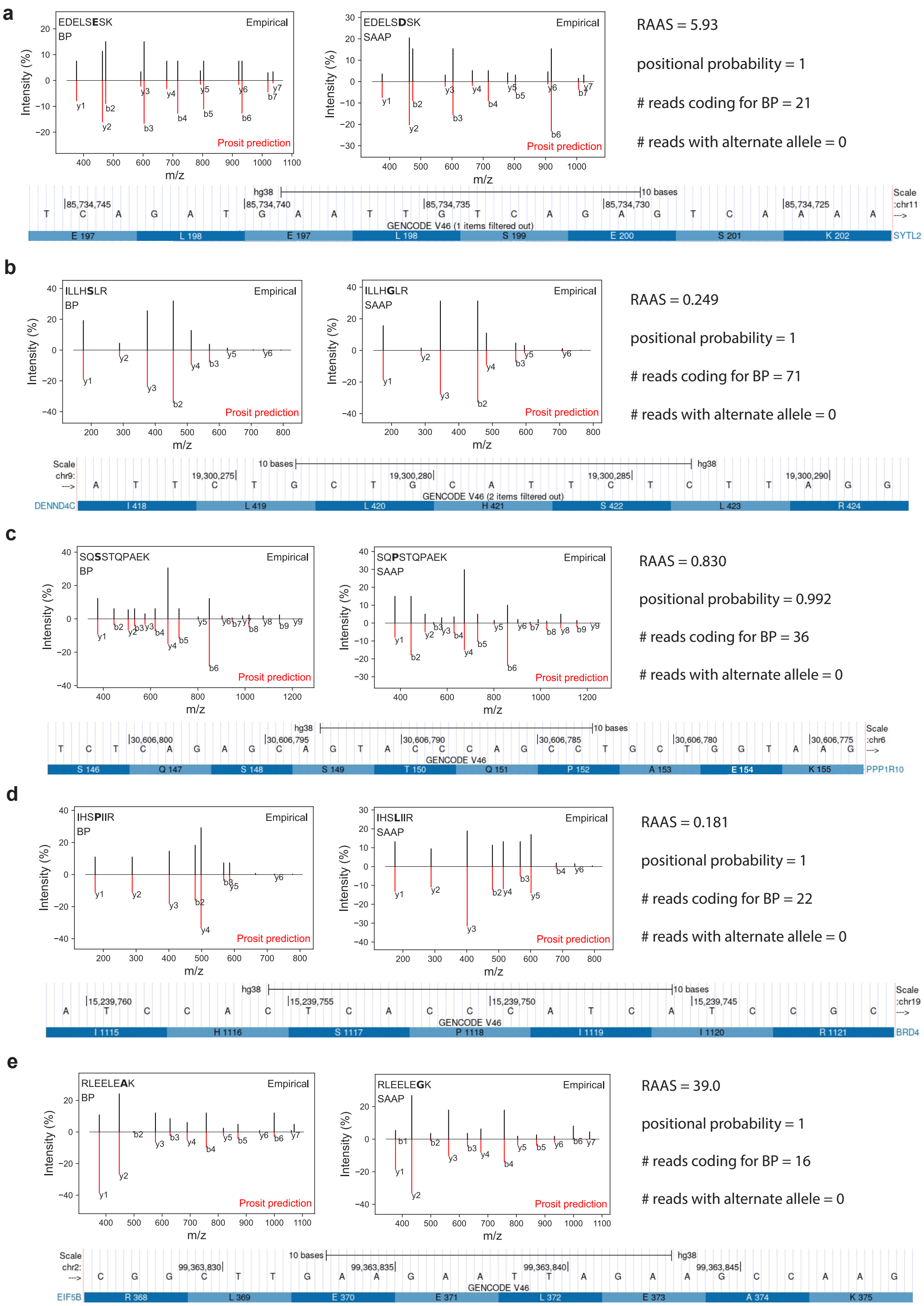


**Supplemental Fig. 2 | RNA-seq and fragment ion evidence for SAAP with high RAAS (part II).** Additional examples of RNA data and mass spectra (TMT-labeled) supporting high RAAS substitutions. All notation is as in Supplemental Fig. 1

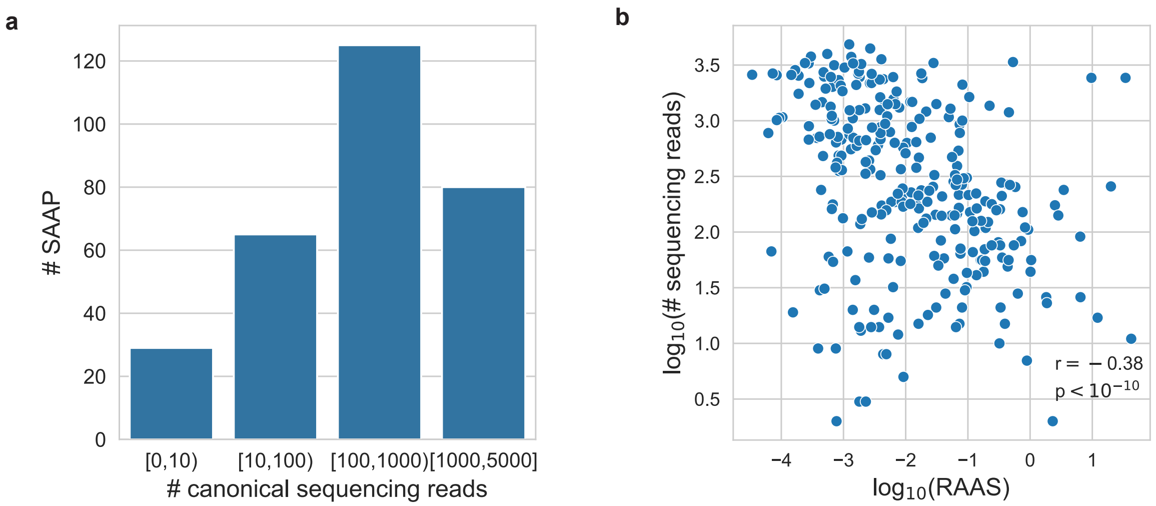


**Supplemental Fig. 3 | Read depth of canonical transcript at substituted peptide locations** (**a**) Number of sequencing reads at each transcript site that encodes for an amino acid for which a substitution was identified. LSCC data only. (**b**) Significant negative correlation between read depth and substitution ratio.

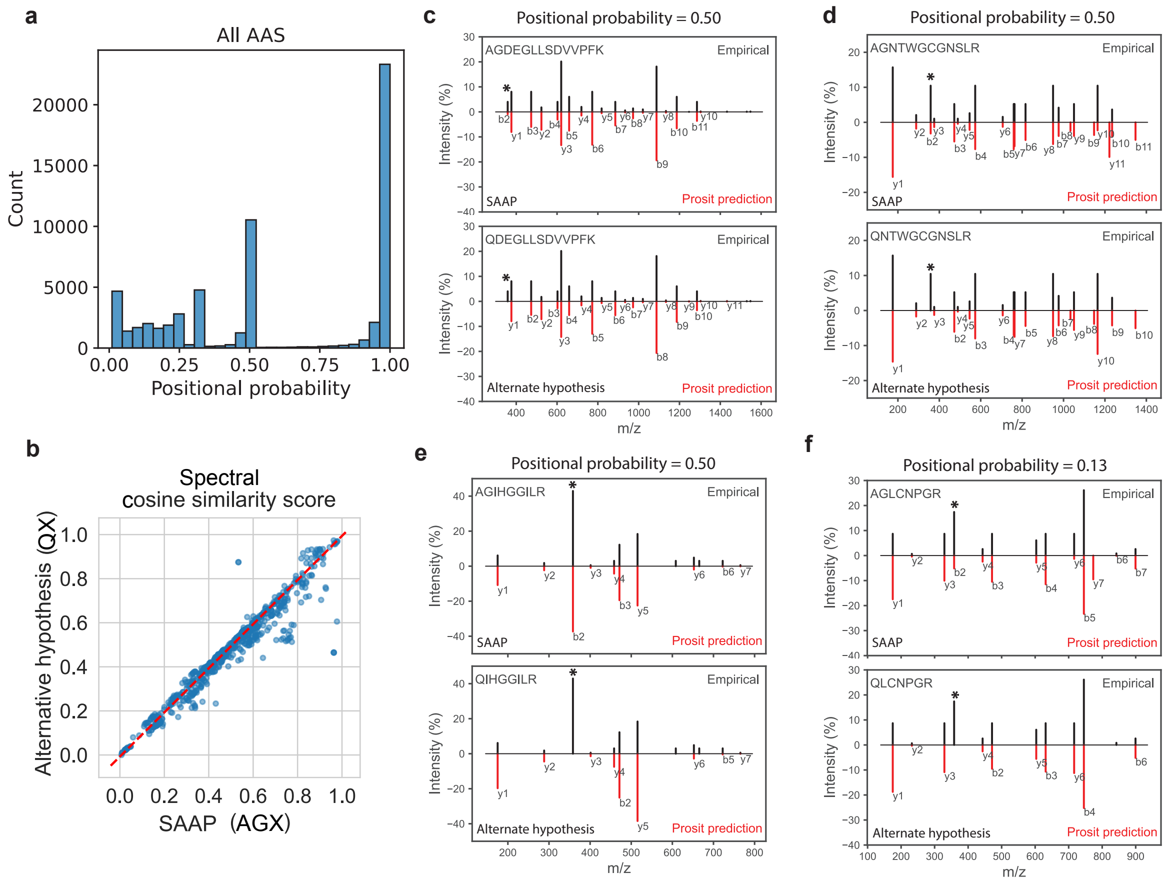


**Supplemental Fig. 4 | Localizing mass shifts and distinguishing between alternative hypotheses for the mass shifts** (**a**) Distribution of the positional probabilities, which quantifies the confidence with which a mass shift can be assigned to an amino acid residue, for all substituted peptides identified by the validation search of the CPTAC and healthy tissues data. (**b**) Cosine similarity score between observed and theoretical spectra for SAAP and alternate peptide sequence suggests that for at least some proposed $Q \to G$ substitutions, the observed spectra better supports the SAAP sequence than the alternate sequence resulting from loss of Ala before Gln. **c-f** Mirror plots of predicted and empirical fragment ion spectra for TMT-labeled SAAP with $Q \to G$ substitutions with uncertain localization of the mass shift. Mirror plots for an alternate hypothesis (alanine cleavage at the N-terminus of the peptide) that is consistent with the mass shift are also shown. The empirical spectra are better represented by the predicted spectra for the $Q \to G$ substituted peptides than for the alternate hypothesis peptides. This conclusion is supported by cosine similarity scores and the presence of peaks (annotated with a star) that match the b2 fragment ion in the spectra predicted for the $Q \to G$ substituted peptides. See Methods for details.

**a**

**Searching protein pull downs for bait proteins with identified SAAPs**

| | |
|---|---|
| Total # baits searched | 25 |
| # baits with detected SAAP generated from bait protein sequence | 18 |
| # unique SAAPs identified | 112 |
| # SAAPs identified mapping to bait | 34 |

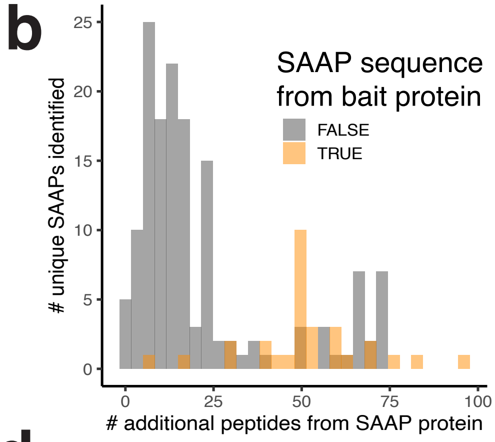


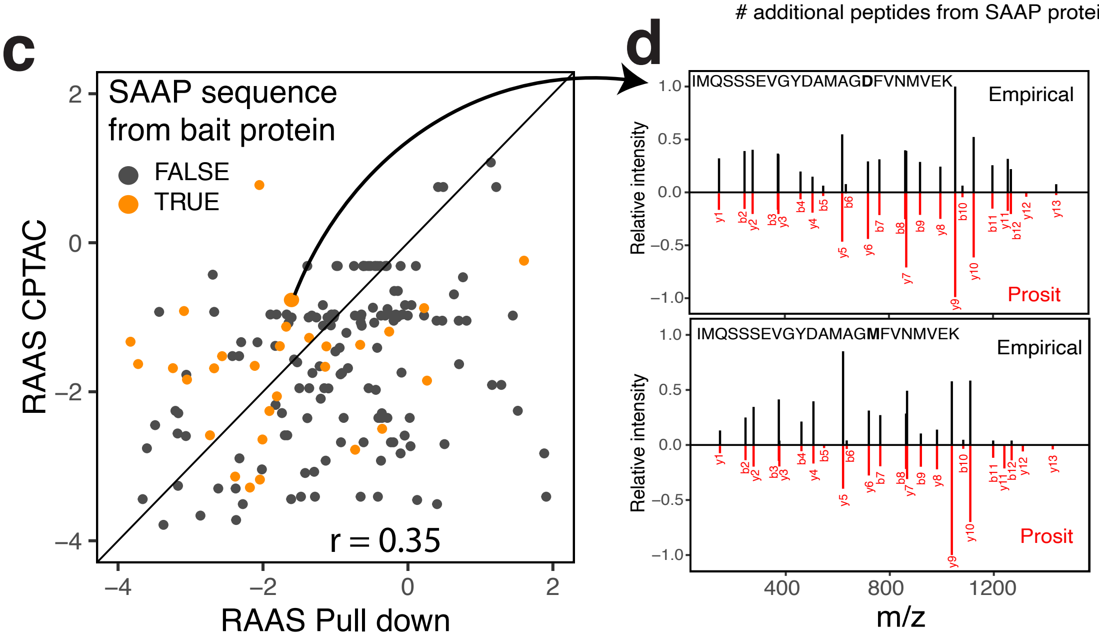


**Supplemental Fig. 5 | SAAP validation from IP-MS experiments** (**a**) A table showing the results from searching 25 different IP-MS experiments for bait proteins with SAAPs identified from CPTAC analysis. (**b**) For each SAAP identified in a given IP-MS experiment, the number of additional identified peptides from the SAAP generating protein are plotted. All but 1 SAAP identified had additional peptides identified. (**c**) The RAAS ratios for identified SAAPs compared to the average precursor RAAS ratios from peptides identified in CPTAC analysis. (**d**) The spectra for a base and substituted peptide pair with prosit predicted spectra.

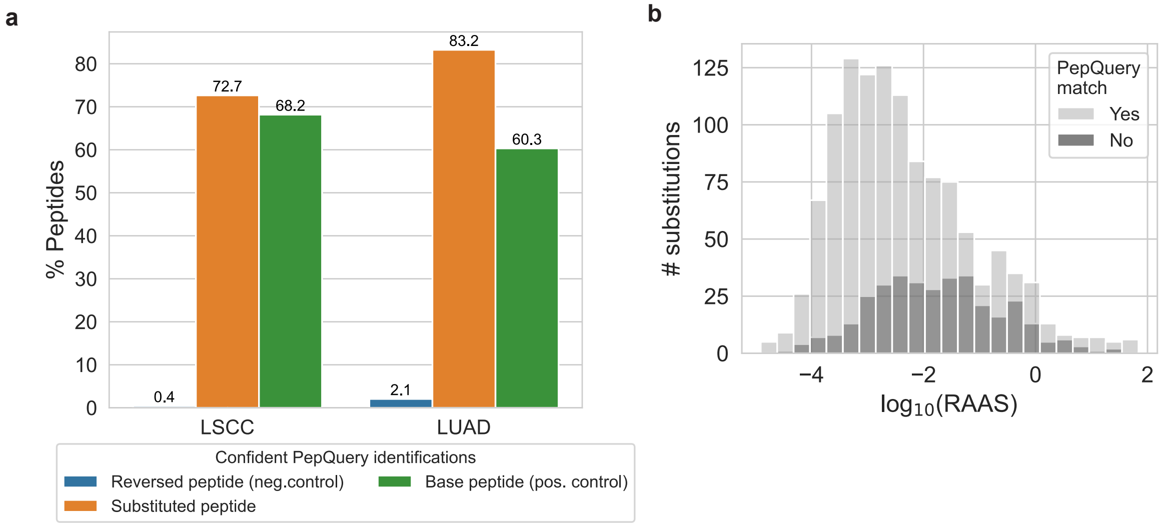


**Supplemental Fig. 6 | SAAP validation with PepQuery** (**a**) Percentage of substituted and control peptides confidently matched to spectra with PepQuery[30]. Encoded peptides are the base peptides corresponding to the substituted peptides. Unrelated peptides are random peptides selected from the dataset to have the same abundance distribution as the substituted peptides. Reversed peptides are the reverse substituted peptide sequences. (**b**) RAAS distributions for substituted peptides stratified by those matched to a spectra with PepQuery and those without a match.

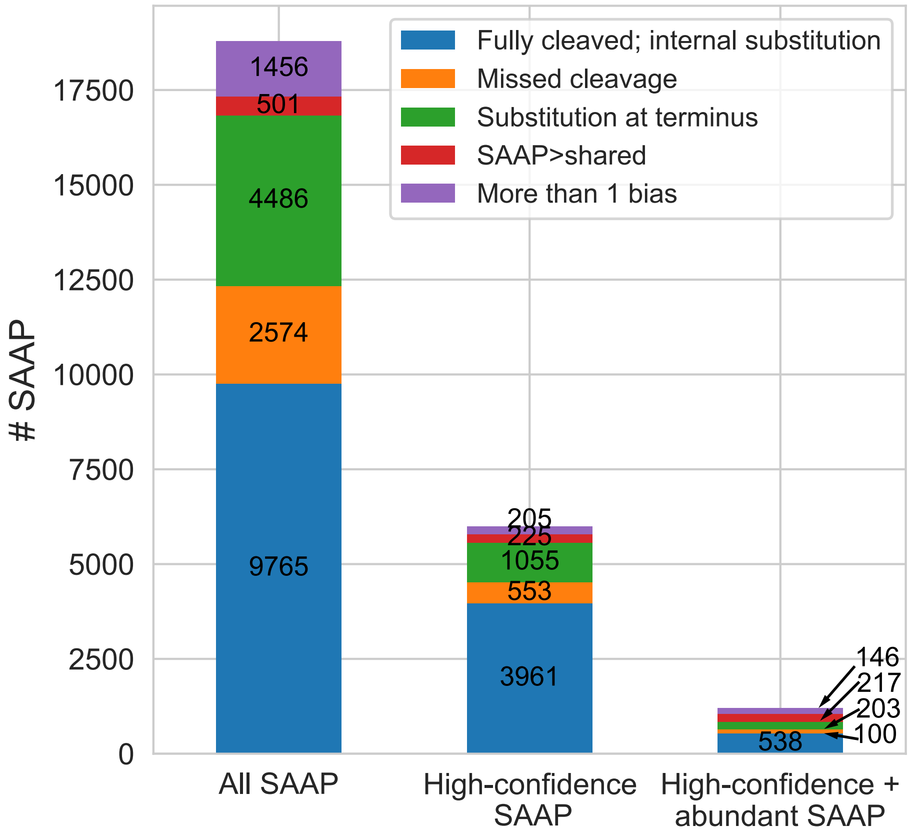


**Supplemental Fig. 7** | Majority of SAAP remain after stringent filtering Filtering of all SAAP, high-confidence (positional probability $\geq 0.9$) and high-abundant (RAAS $\geq 0.1$) SAAP to eliminate those with potential biases leaves the majority of identified substitutions valid for analysis. Biases include missed cleavage, substitution in first or last 2 positions of the peptide, and SAAP that are reported to be more highly abundant than the median of the shared peptides.

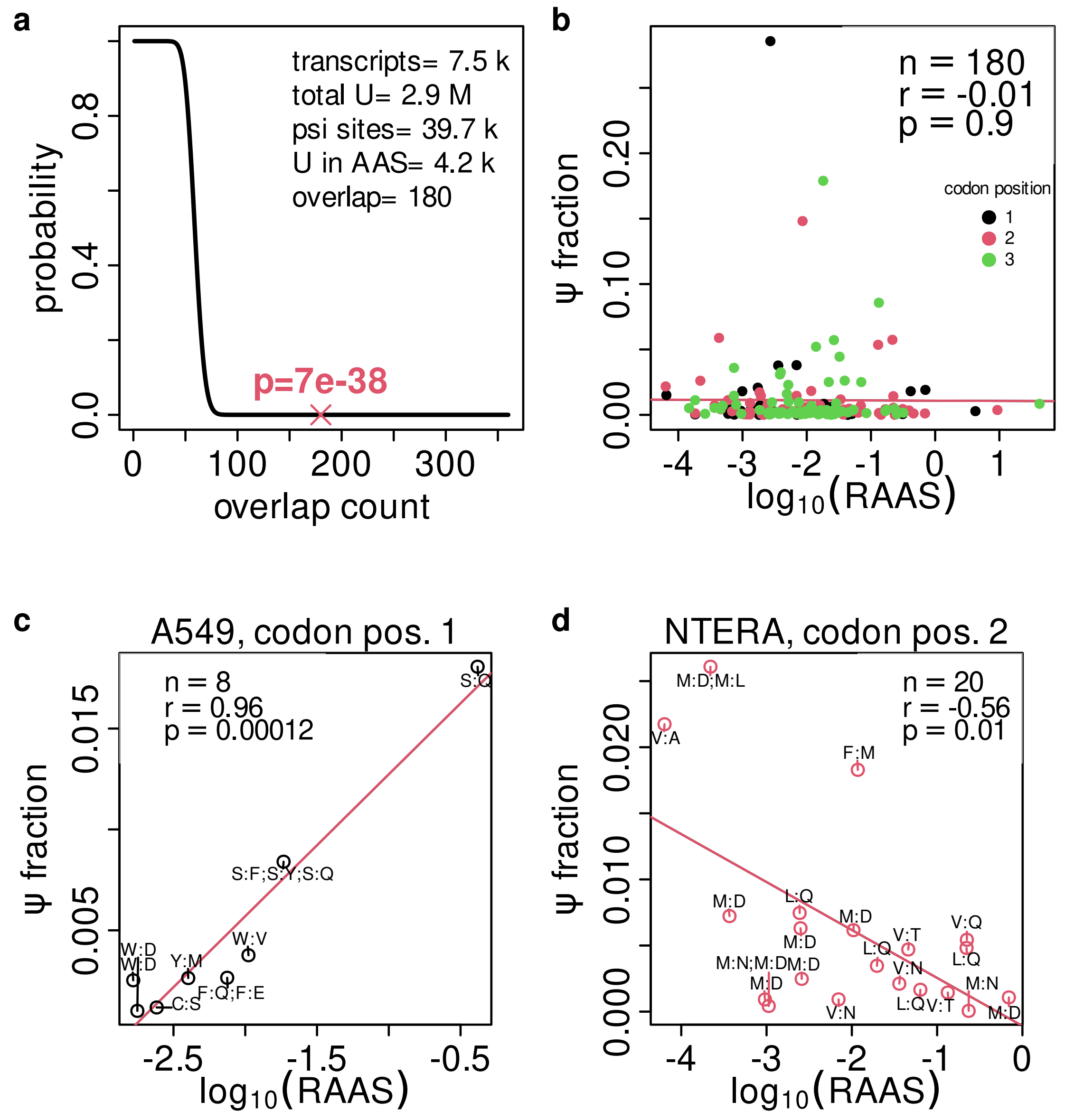


**Supplemental Fig. 8 | Overlap of amino acid substitution with RNA modification sites** (**a**) Modified uracil nucleotides (defined by nanopore sequencing[38]) overlap with codons of amino acid substitution sites. To evaluate the significance of the 180 overlapping sites, we calculated a p-value by a cumulative hypergeometric distribution test (see Methods for details) and here indicate this p-value (red x) in the context of the p-value distribution over different counts of overlaps with otherwise unchanged background numbers. (**b**) The measured fraction of modified U ($\psi$ fraction corresponds to the column `mm.DirectMINUSmm.IVT` in the original data set) is not correlated to the median RAAS of these sites globally. (**c**) The $\psi$ fraction at sites measured in the A549 cell line and overlapping with codon position 1 of amino acid substitution sites is positively correlated to the median RAAS at these sites. (**d**) The $\psi$ fraction at sites measured in the NTERA cell line and overlapping with codon position 2 of amino acid substitution sites is negatively correlated to the median RAAS at these sites.

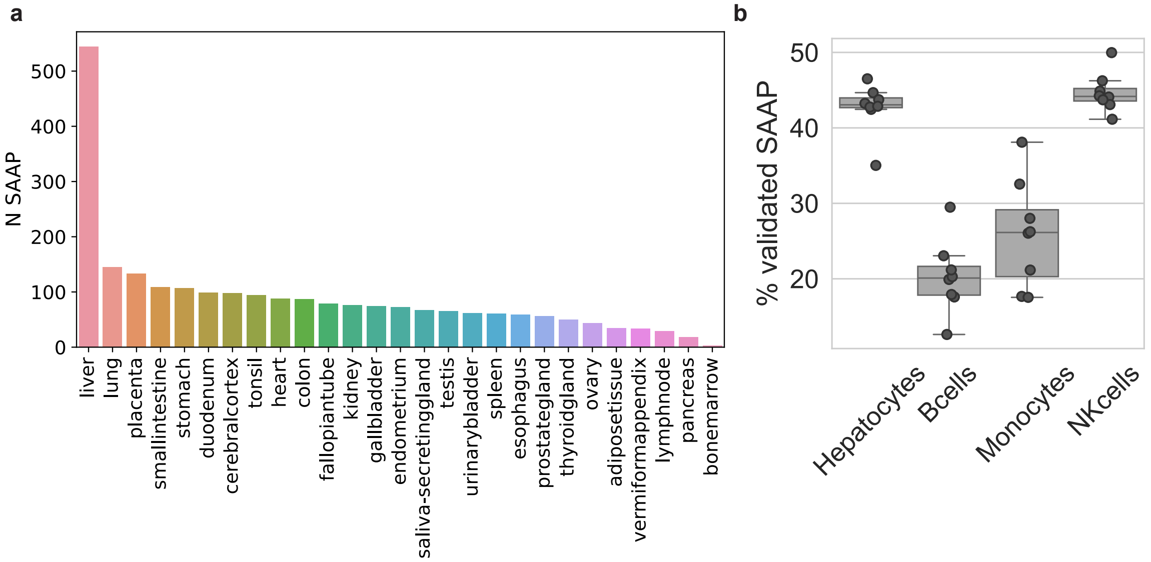


**Supplemental Fig. 9 | SAAP detection and validation in metabolic pulse data** (**a**) SAAP detected in the CPTAC/label-free healthy data analysis that were also detected in the metabolic pulse data from Savtitski, *et al.*[39] are majorly derived from the analysis of healthy liver tissue. (**b**) Percentage of candidate SAAP from dependent peptide search of Savtitski, *et al.*[39] data that were validated by MSFragger.
